# Plasticity in structure and assembly of SARS-CoV-2 nucleocapsid protein

**DOI:** 10.1101/2022.02.08.479556

**Authors:** Huaying Zhao, Ai Nguyen, Di Wu, Yan Li, Sergio A. Hassan, Jiji Chen, Hari Shroff, Grzegorz Piszczek, Peter Schuck

## Abstract

Worldwide SARS-CoV-2 sequencing efforts track emerging mutations in its spike protein, as well as characteristic mutations in other viral proteins. Besides their epidemiological importance, the observed SARS-CoV-2 sequences present an ensemble of viable protein variants, and thereby a source of information on viral protein structure and function. Charting the mutational landscape of the nucleocapsid (N) protein that facilitates viral assembly, we observe variability exceeding that of the spike protein, with more than 86% of residues that can be substituted, on average by 3-4 different amino acids. However, mutations exhibit an uneven distribution that tracks known structural features but also reveals highly protected stretches of unknown function. One of these conserved regions is in the central disordered linker proximal to the N-G215C mutation that has become dominant in the Delta variant, outcompeting G215 variants without further spike or N-protein substitutions. Structural models suggest that the G215C mutation stabilizes conserved transient helices in the disordered linker serving as protein-protein interaction interfaces. Comparing Delta variant N-protein to its ancestral version in biophysical experiments, we find a significantly more compact and less disordered structure. N-G215C exhibits substantially stronger self-association, shifting the unliganded protein from a dimeric to a tetrameric oligomeric state, which leads to enhanced co-assembly with nucleic acids. This suggests that the sequence variability of N-protein is mirrored by high plasticity of N-protein biophysical properties, which we hypothesize can be exploited by SARS-CoV-2 to achieve greater efficiency of viral assembly, and thereby enhanced infectivity.

## Introduction

Two years into the COVID19 pandemic, intense research into the structure and molecular mechanisms of SARS-CoV-2 virus has led to the rapid development and deployment of several types of vaccines (1), neutralizing monoclonal antibodies (2), and small molecules drugs such as ribonucleoside analogs and inhibitors of viral proteases and polymerase (3–5). A persistent concern is viral escape through evolution of therapeutic and immunological targets. Most attention in this regard is devoted to the viral spike protein that facilitates viral entry (6, 7), though recent data additionally point to the importance of viral packaging by the nucleocapsid (N) protein modulating viral loads and thereby infectivity (8–10).

In an unprecedented global effort, several millions of genomes have been sequenced so far and submitted to the Global Initiative on Sharing All Influenza Data (GISAID) to monitor SARS-CoV-2 variants (11) (Fig. 1a). This has provided an invaluable data base for recognizing emerging variants of concern, phylogenetic analyses, and analyses of geographic spread (12–16). The vast majority of the observed mutations are short-lived and ostensibly inconsequential. However, such mutations play a key role in establishing the genetic diversity of RNA viruses, with profound impact on evolutionary dynamics (17–20). In accumulation they define a mutational landscape that is intimately related to stability constraints and structure-function relationships of viral proteins (21, 22). Thus, projecting the ensemble of mutations in reported sequences into the amino acid sequence space and neglecting their origin and relationships, they manifest an exhaustive mutational landscape of viable amino acid substitutions of SARS-CoV-2 proteins, all of which evidently have been successfully proliferating as part of the SARS-CoV-2 species found in patients (16, 23, 24). Unfortunately, the fine-grained interpretation of such mutation data in the biophysical context of protein structure and function is a daunting task, and the spectrum of biophysical properties defined by the ensemble of mutant sequences is largely unclear, although it has been observed that RNA virus proteins generally have more loosely packed cores and intrinsically disordered regions that may provide adaptability (25). Focused analyses of RNA virus mutational landscapes in relation to protein structures have been carried out, for example, for the poliovirus polymerase (21), and the SARS-CoV-2 spike protein receptor binding domain (26, 27).

**Figure 1.**
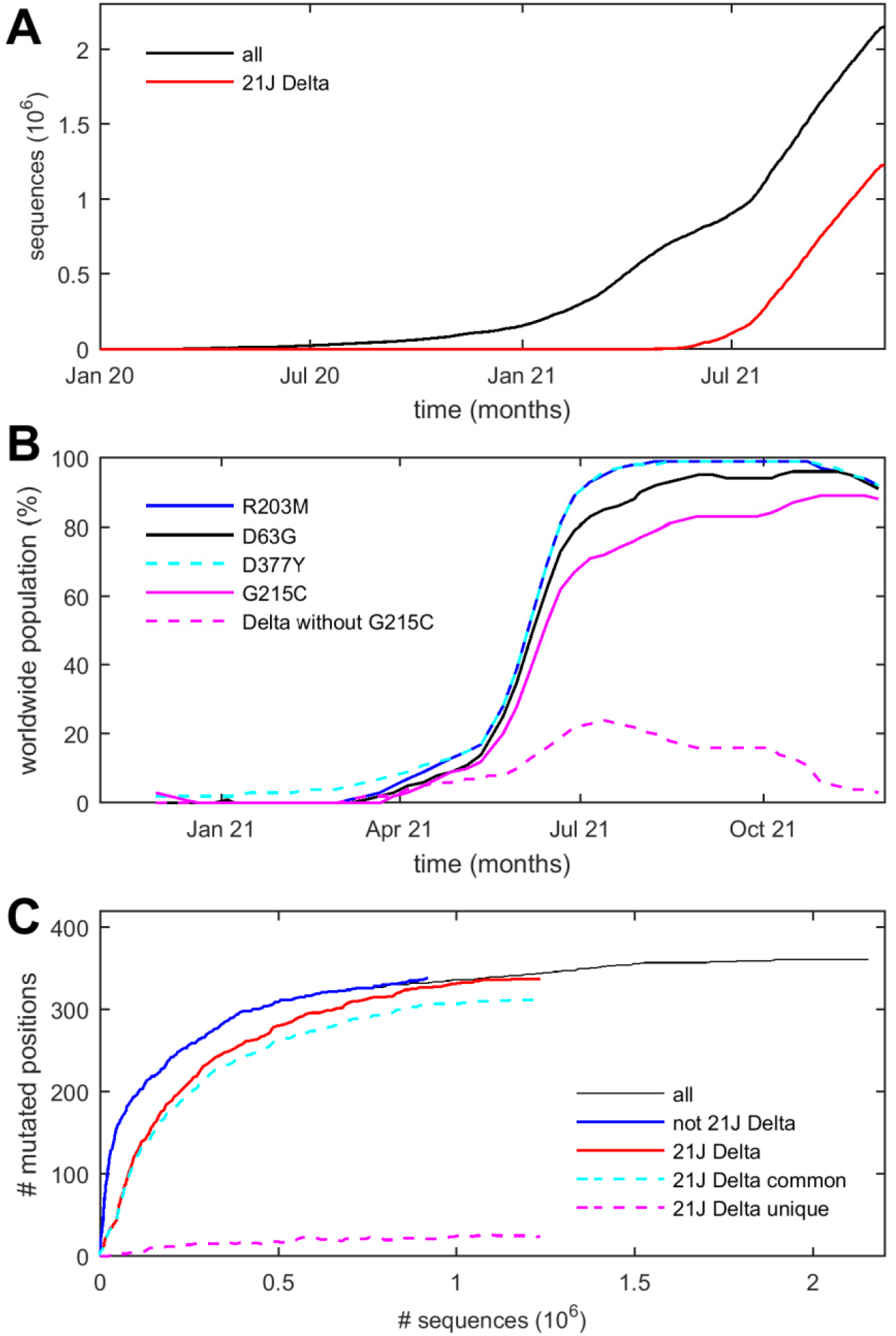
Timeline of SARS-CoV-2 sequences and N-protein mutations. **a** Cumulative number of sequences originating from GISAID and preprocessed by Nextstrain.org. All sequences (black) are shown and those of 21J Delta variant (red). **b** Relative worldwide populations of N-protein sequences exhibiting the characteristic Delta variant mutations D63G, R203M, D377Y, and G215C, assembled from data of Nextstrain.org (13). To highlight the emerging role of G215C, pointed out first by Marchitelli et al. (35) and Stern et al. (34), the dashed line shows the declining contributions of sequences that do not carry the G215C mutation (clades 21A and 21I). **c** Cumulative number of residues at which any substitution was observed versus total sequences. The total (black) is subdivided in those of 21J Delta clade (red) and all others (blue). Also shown are residues in 21J Delta that are common (cyan) or unique (magenta) to this variant.

In the present work we aim to exploit the mutational landscape of the SARS-CoV-2 N protein as a tool to elucidate molecular aspects of the viral assembly, which requires packaging the RNA through an as-of-yet incompletely understood co-assembly mechanism with N protein into well-defined ribonucleoprotein particles (28, 29). Of particular interest with regard to viral assembly is the SARS-CoV-2 variant B.1.617.2 (Delta) that in 2021 has rapidly outperformed all previous variants (30) (Fig. 1a), exhibiting reduced incubation time and significantly higher viral load in infected patients (31–33), in one study up to a 1000-fold higher compared to the original lineage (31). Delta variant mutations in N-protein include D63G, R203M, D377Y, and in different clades additionally R385K and G215C, respectively (34). Among those, mutation R203M was shown to significantly increase replication (8), but is similar to analogous N-protein mutations in all other SARS-CoV-2 variants of concern (34) including the current Omicron variant. Conspicuously, the Delta variant containing the G215C mutation arose without accompanying changes in the spike protein (Nextstrain clade 21J) and has dramatically outperformed and virtually displaced other Delta variant clades, assuming worldwide dominance in 2021 (34, 35) (Fig. 1b). This warrants a detailed study of the impact of the G215C mutation on protein structure and function.

In the present work, we observe a highly conserved region that reveals a possible role of the G215C mutation in enhancing interactions critical for assembly. To examine molecular mechanisms in detail, we combine biophysical characterization of protein size, shape, structure, and protein interactions with structural models from molecular dynamics simulations. In comparison with biophysical properties of the ancestral N-protein, the G215C mutant displays significant differences in secondary structure, self-association and co-assembly with nucleic acid, revealing a plasticity of protein biophysical properties that mirrors its remarkable sequence variability.

## Results

### The Mutational Landscape of SARS-CoV-2 N-protein Reflects Structural Features

Among 2.49 million SARS-CoV-2 sequences uploaded to GISAID and subsequently preprocessed at Nextstrain.org (13) as of November 29, 2021, we find 8.2 million instances of amino acid changes in the N-protein. Outside the consensus mutations associated with different clades (see below), each sequence exhibits on average only 1.3 additional mutations in N-protein. From inspection of phylogenetic trees at Nextstrain.org, a majority of mutations arose multiple times independently, but usually persisted only briefly. Despite their sparsity, when aggregated over 10^6^ sequences, the mutation data is highly redundant, describing 1,264 distinct mutations observed a median of 70 times, assembled in different combinations within 24,982 distinct N-protein sequences (Fig. 2). Mutations occurred for 362 out of 419 residues of the N-protein, each on average allowing 3.5 different substitutions. In range and depth of variability, N-protein exceeds all other structural SARS-CoV-2 proteins as well as ORF1ab (SI Appendix Table S1). Notably, this plasticity of the amino acid sequence does not extend to most of the 37 positions strictly conserved across related betacoronaviruses, 30 of which exhibited no or only conservative substitutions in SARS-CoV-2 (Fig. 2).

**Figure. 2.**
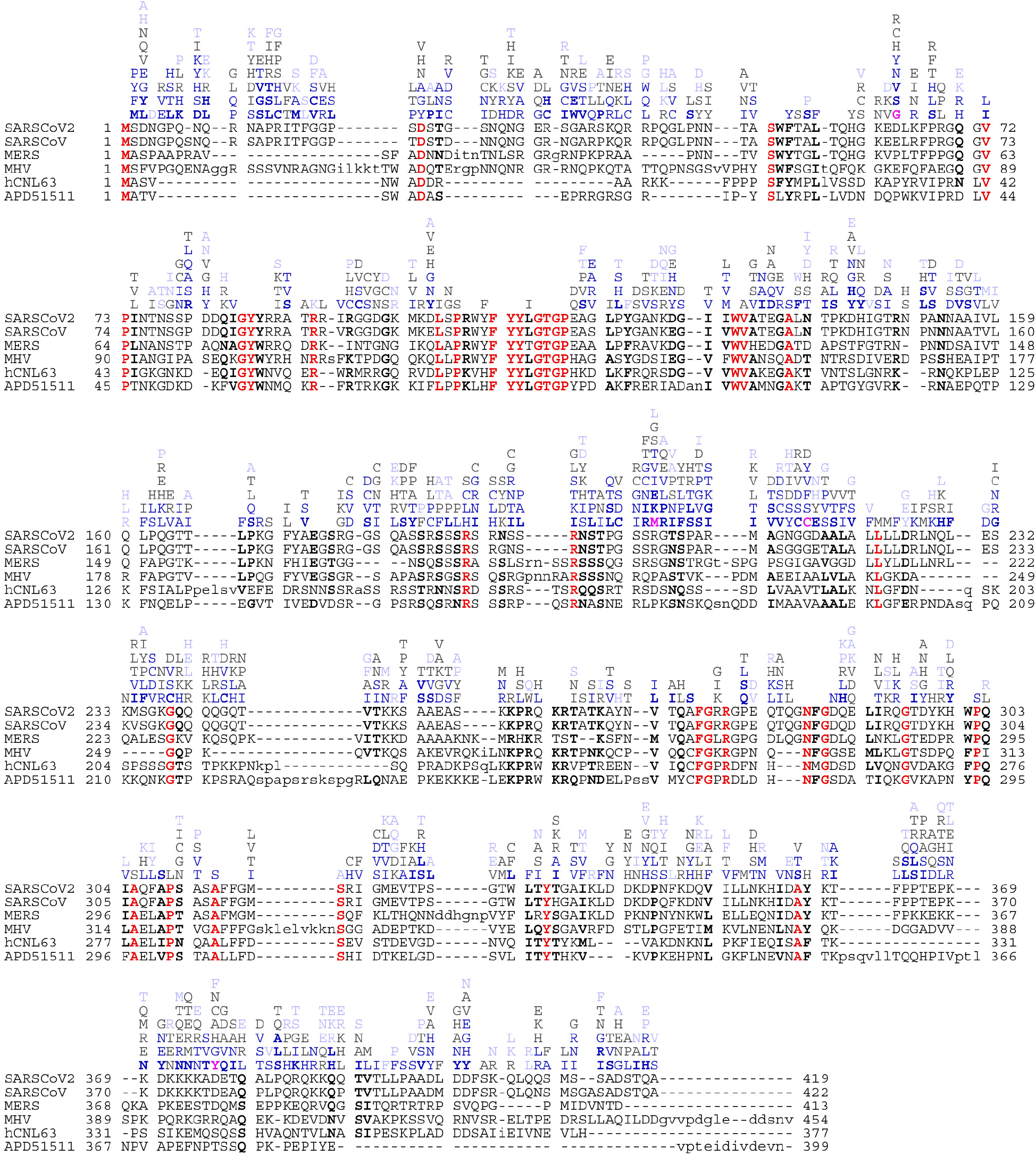
Observed mutations of N-protein in comparison with sequence alignment of related betacoronaviruses. The alignment shows Wuhan-Hu-1 SARS-CoV-2 (P0DTC9.1), SARS-CoV-1 (P59595.1), MERS (YP_009047211.1), murine hepatitis virus (NP_045302.1), human coronavirus NL63 (Q6Q1R8.1), and the 229E-related bat coronavirus APD51511.1. Conserved identical residues are highlighted in bold red, and conserved similar residues in bold black. Above the aligned sequences are observed distinct mutations of SARS-CoV-2 observed among 1.69 million sequences uploaded to GISAID since beginning of the pandemic, ordered by frequency < 10–20 (light blue), 20–100 (gray), 100–1,000 (blue), and >1,000 times (bold blue).

To assess to what extent the observed sequence space has exhausted the range of possible substitutions, Fig. 1c shows the scope of residues at which mutations have been observed as a function of total sequences. (Because we assume most sequence fluctuations to be stochastic independent events, the cumulative number of sequences is serving here as a scaled surrogate for time to compensate for vastly different sequence deposition rates with time.) The approach of an asymptotic limit can be discerned, with a second phase of changes in emerging Delta variant sequences; these, too, approach a limit. This suggests that the observed mutation landscape approximates an equilibrium, while hinting at some changes associated with the Delta variant. To further analyze the time course of observed mutations, Fig. 3e presents the temporal evolution of the mutation frequency observed at each position. Insofar as the propagation of mutations in each N-protein position are driven by adventitious spreading events, or are bystander of improvements in fitness to other viral proteins, variation along the ordinate (time/sequence axis) in Fig. 3e is produced. By contrast, constant mutation frequencies in individual residues (constant color along vertical line in Fig. 3e) must reflect intrinsic molecular properties. Remarkably, for the overwhelming majority of positions across the N-protein, the observed mutation frequency is largely constant with minor stochastic modulations; this becomes even clearer when subdividing sequences between Delta 21J clades and others pre-dating 21J (SI Appendix Fig. S1). However, from the observed time-course of ancestral SARS-CoV-2 and Delta 21J mutations, it did require on the order of 10^5^ sequences to approximate the mutational landscape (SI Appendix Fig. S2).

**Figure 3.**
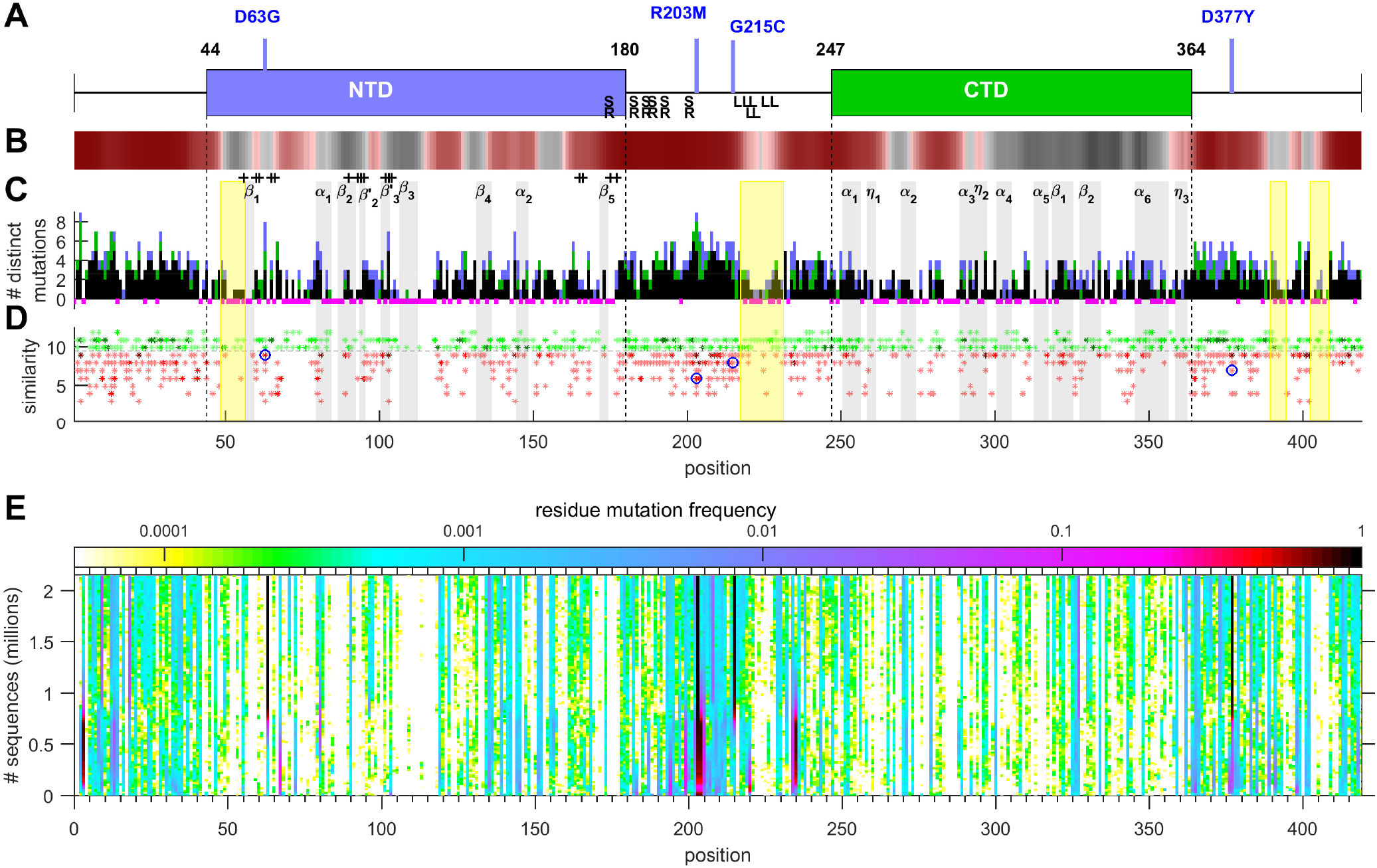
Structural organization of N-protein and range and frequency of mutations. **a** Domain organization of N-terminal and C-terminal folded domains with highlighted mutations characteristic for Delta variant SARS-CoV-2. **b** Propensity of residues to promote LLPS, showing in red droplet-promoting regions with pDP >0.6. **c** Histogram of the number of distinct mutations of each residue. The histogram bars are subdivided to show the number of distinct mutations common to 21J Delta and non-21J Delta species (black), those that have only occurred in non-21J Delta (green) and those only observed in 21J Delta species (blue) (see SI Appendix Fig. S4). Magenta highlights below the abscissa indicate positions with no or only conservative substitutions. Grey vertical patches indicate regions of secondary structure identified by Dinesh and colleagues from NMR of the NTD and by Zinzula et al. from x-ray crystallography of the CTD, and + signs indicate positions with NMR chemical shifts upon nucleic acid binding in the NTD (43, 44). Yellow patches indicate regions of unknown function that appear conserved from mutations. **d** Mutations scored for physicochemical similarity with conservative substitutions green and non-conservative red. Multiple instances of the same score are depicted as darker shade. Blue circles highlight the four Delta variant mutations. **e** Frequency of mutations *vs.* total sequence number. To visualize the temporal fluctuations in the rate of accumulated mutations, the daily number of any mutation in each position relative to the daily number of new sequences, convoluted across 10^4^ sequences, is plotted against the total accumulated sequences (as scale of time) and color-coded according to the relative rate of observing a mutation. Thus, structure in the vertical direction reflects adventitious spreading events and mutational drifts, superimposed to constant baseline mutation fitness dictated by molecular properties. At ∼10^6^ sequences, a slight change in pattern may be discerned, coinciding with the rise of the Delta variant. Mutation frequency plots separately for Delta 21J and preceding variants are in SI Appendix Fig. S1; their initial spread in the mutational landscape in shown SI Appendix Fig. S2.

We can better examine the significance of the mutations in more detail in the context of the N-protein structural organization and its assembly function. Briefly, N-protein is dimeric, with each chain comprised of a C-terminal dimerization domain (CTD) and an N-terminal nucleic acid (NA) binding domain (NTD), both flanked and linked by long intrinsically disordered arms (Fig. 3a) (36). Both domains and linker contribute to promiscuous NA binding. NA binding induces a more ordered conformation that allows dimer-dimer interactions, which, in turn, and in concert with scaffolding on NA, lead to multimeric co-assemblies (37). NA binding also promotes liquid-liquid phase separation (LLPS), and the highly concentrated co-condensates of N-protein and NA allow the formation of ribonucleoprotein particles (38–41). N-protein also interacts with the viral M-protein, which appears to play a role in promotion of N-protein condensates, in anchoring ribonucleoprotein particles to the viral membrane, and in the recognition of viral RNA (39, 42).

Overall, the number of distinct mutations tracks well with the folded domains, which have a more restricted range of viable mutations (Fig. 3c). Residues close to the secondary structure elements previously revealed by NMR spectroscopy and x-ray crystallography are mostly either completely protected from mutations or highly conserved, as are the majority of NTD residues likely in contact with nucleic acids, as inferred from NMR chemical shifts (43, 44). Conversely, regions in the linker that score high for disorder and LLPS propensity (Fig. 3b) exhibit the largest clusters of mutations. When examining the nature of the replacements through a scoring for physicochemical properties, such as polarity, hydrophobicity, and secondary structure propensity, we can discern a majority of conservative substitutions in the folded domains, and greatest variety of residues with altered physicochemical properties in the disordered regions (Fig. 3d).

New features appear in this mutational landscape in the disordered linkers and arms. The linker is critical to RNA-mediated phase separation (39) and was early identified as a region of high sequence variability (14). The large coverage of the mutational landscape now available allows identification of constrained islands within the disordered regions. These include positions 49–56 proximal to the first sheet in the NTD, the highly conserved leucine-rich sequence 218–231, as well as two stretches in the C-terminal arm (C-arm) at 390–394 and 403–408 (yellow highlights in Fig. 3c, d). Interestingly, these regions also approximately correspond to distinct stretches of amino acids with low disorder score. Based on their apparent conservation, these regions likely endow N-protein with critical functions.

The disordered C-arm of N-protein is thought to play a critical role in the interaction with viral M-protein and the packaging signal, based on studies with corresponding regions of other coronaviruses (42). The observed conserved stretches in the mutational landscape coincide with helices (383–396 and 402–415) previously observed in molecular dynamics (MD) simulations (38). In further support, we find structural prediction displays transient helices in this region, in the AlphaFold2 result spanning residues 400–410 (SI Appendix Fig. S3). The relatively low confidence score is expected given the intrinsic disorder of the C-arm, but nonetheless suggests a propensity for transient helix formation. Structure predictions of segment Q390-A419 by other methods (I-TASSER and Phyre2) also suggest helical content, with the highest score for the segment 400–408.

For the protected leucine-rich sequence 218–231, a potential role arises from its location within the linker region 210–246 found to be essential for RNA-mediated LLPS (39). The conserved island is also overlapping with the locus of a previously identified nuclear export signal 224–230 (45). Incidentally, it overlaps with the peptide 222–230 that is a binder for HLA-A02:01 and immunogenic (46). Cubuk et al. reported transient helices in the leucine-rich region in MD simulations, proposed to provide oligomerization interfaces (38). Using structure-prediction tools, we confirm the presence of a helical segment spanning residues 215–235 (SI Appendix Fig. S3), and, as further described below, and find evidence for its role in protein oligomerization and co-assembly with NA.

On the N-terminal half of the disordered linker, the SR-rich region 176–206 has been a locus of particular interest due to the cluster of charged residues and phosphorylation sites. Their phosphorylation state is thought to regulate N-protein functions (36, 39, 40, 47) and interaction with the viral NSP3 protein (47) and host proteins such as glycogen synthase kinase-3 (48, 49), CDK-1 (40), and 14-3-3 proteins (50). Focusing on the 14 serine residues in this stretch, the mutational landscape shows that, with the exception of completely conserved S176, all other phosphorylation sites can be substituted, and new ones can be introduced. Overall, 14.3% of all sequences exhibit changes in the serine pattern; however, there is a significant anti-correlation to maintain a total number of 13 or more serines in this region (**Table S2**). Thus, it appears that except for S176, there is redundancy and flexibility in the phosphorylation sites, but with a constraint to maintain their local density.

Within this SR-rich region, the R203 mutation was noted early in the pandemic and remained common to all variants of interest (34, 51, 52). The earlier R203K/G204R was shown experimentally to enhance the ability of N-protein to form condensates (6), and R203M – prevalent in the Delta variant – was recently reported to enhance viral replication (8).

The most recent mutation G215C is located in the linker between the SR-rich and leucine rich regions. It has arisen in the Delta 21J clade during the last several months. Accompanied solely by mutations in ORF1ab and ORF7b, it quickly outcompeted any variants not containing G215C to assume worldwide dominance (Fig. 1b). As of November 29, 2021, 49.6 % of sequences in the data base contain the G215C mutation, but these describe only 34.9% of unique sequences (8,720), consistent with the shorter period of time since the 21J clade has emerged. Nevertheless, when examining to what extent these have already expanded to reproduce the mutational landscape, we find 97.2% of instances of N mutations occur in positions that show mutations in both 21J and non-21J clades (Fig. 3c, see SI Appendix Fig. S4 for a detailed list). Incomplete overlap can be discerned mostly among rare substitutions and in highly disordered regions. Quantitatively, 14.0% of distinct mutations in non-21J (pre-Delta) species have not yet been observed in the 21J clade, potentially due to still incomplete coverage. Interestingly, however, 21.8% of distinct substitutions in the 21J clade were not previously observed in non-21J clades, suggesting evolution of biophysical properties (Fig. 1c).

### N-protein mutants display altered secondary structure and assembly properties

To study the impact of the mutation on structure and assembly function of N-protein, we compare the biophysical properties of select N-protein mutants derived from the Delta variant with those of the ancestral Wuhan-Hu-1 N-protein (Nref) *in vitro*. First, we examined the coarse-grained size and shape using sedimentation velocity analytical ultracentrifugation (SV) and dynamic light scattering (DLS). As shown previously (37), when expressed in *E*. *coli* and purified to remove NA, Nref forms non-covalent 4.1 S dimers with a Stokes radius of 5.9 nm. Its translational frictional ratio of 1.82 indicates a highly extended hydrodynamic shape as a result of significant disorder. The dimers are linked tightly at the dimerization interface in the CTD with *K*_*D*_ < 10 nM, but show only ultra-weak further self-association (*K*_*D*_ = 760 μM).

An overlay of sedimentation coefficient distributions of the N:D63G and N:G215C mutants, as well as the quadruple N:D63G,R203M,G215C,D377Y mutant reflecting the full set of canonical mutations in 21J Delta variant is shown in Fig. 4. While N:D63G sediments similar to Nref, indicating no change in size, shape, or self-association properties, both N:G215C and the quadruple mutant sediment much faster, at a rate that demonstrates the formation of tetramers at low micromolar concentrations. To comprehensively examine the altered state we focus on N:G215C, in light of its unique epidemiological impact, and to enable clear structural attribution of changes to a single residue substitution.

**Figure 4.**
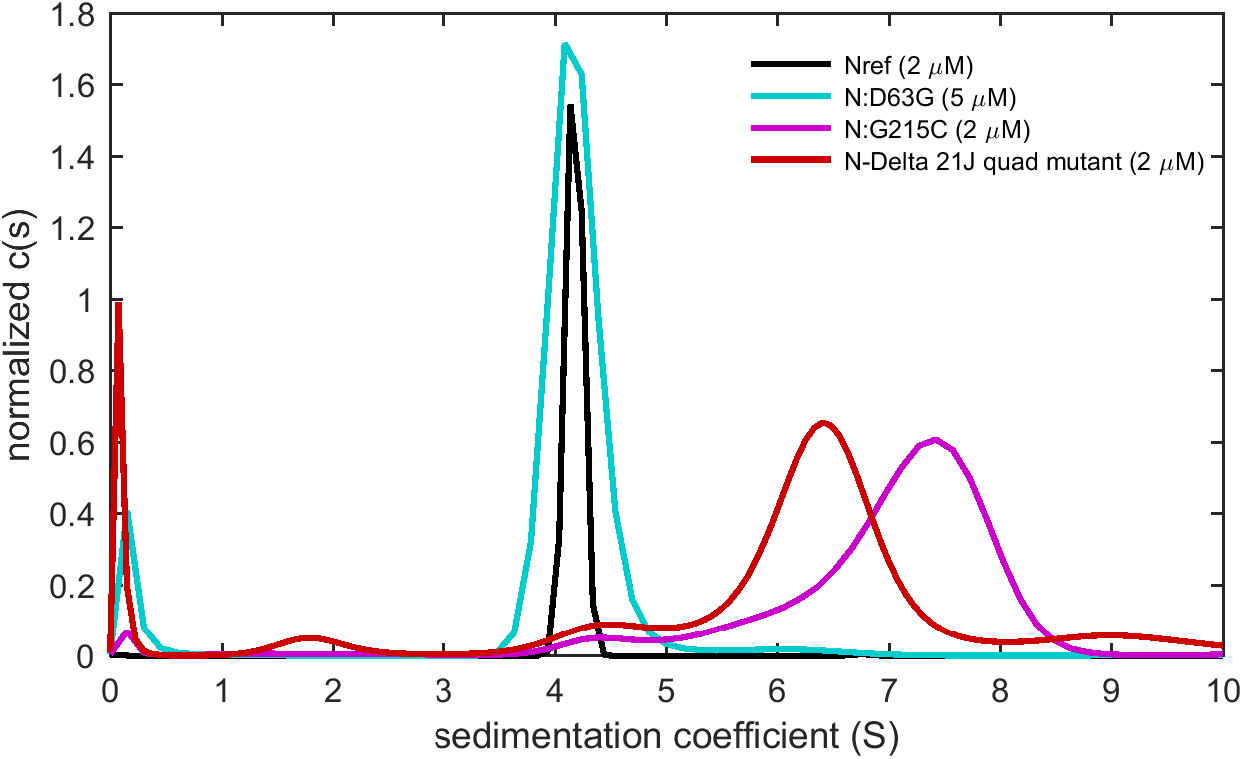
Self-association properties differ for N-protein mutants. Sedimentation coefficient distributions are shown for ancestral protein (Nref; black), the N:D63G mutatnt (cyan) the N:G215C mutant (red), and the quadruple mutant N:D63G,R203M,G215C,D377Y that reflects the set of mutations common to the 21J Delta clade (red).

Compared to the ancestral Nref, N:G215C sediments much faster at 7.3 S (Fig. 5a), with a lower hydrodynamic frictional ratio of 1.58, in a relatively more compact, tetrameric solution state with a Stokes radius of 7.09 nm (Fig. 5c). Electrospray mass spectrometry (ESI-MS) shows an intact mass of 93,792 Da, consistent with the expected value for a His-tagged dimer. Furthermore, LC-MS/MS of tryptic digests shows a peptide consistent with a species composed of two tryptic peptides crosslinked by disulfide bond at 215C. Finally, we used mass photometry (MP) to obtain an independent direct measurement of the molecular weight distribution of N:G215C in solution (Fig. 5d). Different from ESI-MS, this method leaves the non-covalent dimerization at the CTD intact. Accordingly, a majority peak can be discerned approximately at the tetramer mass. In summary, N:G215C forms a tightly bound, compact tetramer *via* disulfide crosslinks of non-covalent dimers.

**Figure 5.**
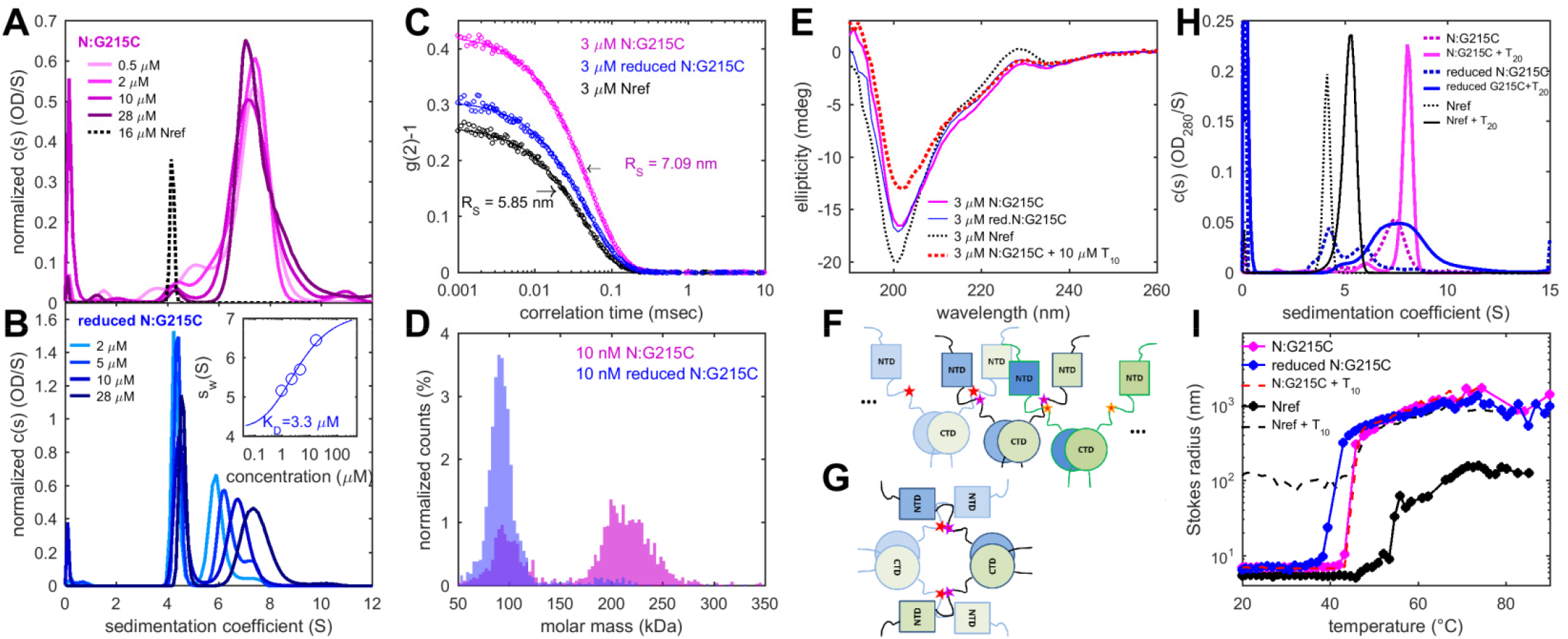
Solution structure and higher-order assembly properties of N:G215C protein. **a** Sedimentation coefficient distributions of N:G215C at a range of concentrations. For comparison, dimeric Nref is shown as dotted line. **b** Sedimentation data in reducing conditions after incubation in working buffer supplemented with 1 mM TCEP. The inset shows the concentration-dependence of the weight-average sedimentation coefficient (circles) and best-fit isotherm model (line) leading to a K_D_ for dimer-tetramer self-association of 3.3 μM. **c** Autocorrelation function in DLS (circles) and best-fit single-species model with R_S_ = 7.09 nm (lines). **d** Determination of mass distribution by MP. Shown are histograms of the masses associated with single molecule surface adsorption events of N:G215C in working buffer (magenta) and in reducing conditions (blue) after incubation with 10 mM DTT. MP is restricted to low nM concentrations, leading to partial dissociation of the CTD dimerization interface. This can be discerned from the minority dimer peak under non-reducing conditions. Due to the lower mass limit of MP of 50 kDa, free N-protein monomer cannot be measured. **e** Secondary structure content of Nref and N:G215C at different conditions by circular dichroism, and conformational changes of N:G215C in presence of T_10_. Spectra for reduced and non-reduced N:G215C virtually overlap. **f, g** Cartoons of N-protein assembly with different chains of CTD-linked dimers (CTD circles, NTD squares, blue and green depicting dimer chains, brightness scale distinguishes different dimers) are forming inter-dimer (**f**) or intra-tetramer (**g**) cross-links at 215C (stars). **h** Sedimentation coefficient distributions of 2 μM N-protein in the presence of 1 μM oligonucleotide T_20_ which can bridge N-protein dimers. For reference, T_20_ binding to Nref leads to weak dimer self-association at these low concentrations (black). For N:G215C (magenta) which is constitutively tetrameric, binding of T_20_ causes an increased sedimentation velocity consistent with the additional bound mass. Reduced N:G215C (blue, with 10 mM DTT) is constitutively in a dimer-tetramer self-association equilibrium, which T_20_ binding shifts strongly to tetramers. For both disulfide-linked and reduced N:G215C weak higher-order co-assembly of the tetramer/T_20_ complex cannot be excluded. **i** Promotion of particle growth at higher concentrations and higher temperature. Shown are the average Stokes radii measured by DLS for 3 μM N:G215C in standard (magenta) and reducing conditions (blue, with 10 mM DTT), and, for comparison, 3 μM NRef in standard conditions (black). Also shown is 3 μM N:G215C in presence of 10 μM T_10_ oligonucleotide (dashed red), as well as 3 μM Nref with 10 μM T_10_ (dashed black).

When disrupting disulfide bonds in reducing conditions, as expected, the intact mass obtained from ESI-MS was 46,897 Da, consistent with the mass of the monomer. By MP, the mass distribution exhibits a peak at the dimer molecular weight (Fig. 5d), consistent with significant non-covalent CTD dimerization similar to Nref. However, at the much higher μM concentrations in SV, reduced N:G215C shows much different behavior (Fig. 5b): In addition to a major peak at 4.1 S for the dimer, a faster sedimenting population with distinct concentration dependence can be discerned. This is characteristic of further self-association in rapid association/dissociation exchange (53). An isotherm of weigh-average *s*-values can be modeled as a dimer-tetramer association step with a best-fit dimer *K*_*D*_ of 3.3 μM (Fig. 5b inset). This is ∼200-fold stronger than previously measured for Nref. Thus, the G215C mutation induces conformational alterations that create, or significantly enhance, a non-covalent dimer-dimer protein interaction interface outside the CTD, even in the absence of covalent disulfide bonds.

The possibility of covalent dimerization of CTD-linked dimers poses a question regarding the quaternary structure, since multivalent dimers could potentially self-assemble into wide range of higher-order oligomers (Fig. 5f). Such structures would be detected with high sensitivity both in SV and DLS but are virtually absent in our data. Instead, as supported by reduction and re-oxidation experiments (SI Appendix Fig. S5), measurement of free sulfhydryls (SI Appendix Fig. S6), and the measured compact hydrodynamic shape (Fig. 5a, c), we propose a more compact configuration with two intra-tetramer crosslinks (Fig. 5g). Unfortunately, published cryo-ET structures do not yet allow unambiguous structural assignment of N-protein configuration in the ribonucleoprotein particles (29).

To complement the study of coarse-grained aspects of N-protein size and shape, we examined the secondary structure content of N:G215C by circular dichroism spectroscopy (CD). Whereas the spectrum of Nref is dominated by a large negative ellipticity at 200 nm that is characteristic for disordered chains, N:G215C shows much reduced negative 200 nm signal and instead stronger ellipticity in the range 220-230 nm typical for helical structures (Fig. 5e). Such diminished disorder is consistent with the more compact hydrodynamic shape of N:G215C. Virtually identical spectra were obtained in reducing conditions, and across a concentration range populating different fractions of dimers and tetramers.

A key step in assembly is the interaction between NA and N-protein. We previously probed consequences of NA binding on N-protein interactions by studying N-protein liganded with short oligonucleotides (37). Up to a decanucleotide T_10_, a length that spans the binding grove of the NTD (43), we observed similar binding affinities of NA for N:G215C as previously determined for Nref (SI Appendix Fig. S7). Likewise, virtually unaltered is a reduction in disorder by CD spectroscopy when liganded by T_10_ (Fig. 5e), as well as a NA binding-related shift to greater thermal stability of the folded domains observed by differential scanning fluorometry (SI Appendix Fig. S8). Apparently, elementary features of NA binding are not substantially affected by the G215C mutation.

A much different picture arises when binding the longer oligonucleotides T_20_. These spatially extend beyond a single NTD domain and can bridge between two N-protein dimers (37). For Nref, they promote formation of tetramers and higher oligomers at low micromolar protein concentrations (37). In the case of N:G215C, we find this co-assembly significantly augmented. While this may be expected for disulfide-linked N:G215C tetramer, even in the reduced conditions N:G215C exhibits significantly stronger hetero-oligomerization (Fig. 5h). This shows cooperativity or an avidity advantage of N:G215C in the earliest steps of assembly with NA, presumably due to its ability to constitutively tetramerize.

Finally, we compare aspects of higher-order assembly and LLPS. LLPS depends on highly multi-valent, weak interactions (54), such as transient aromatic side-chain and backbone interactions of disordered chains (55). These can be expected to differ from protein/NA and protein-protein interactions that stabilize the discrete oligomeric co-assemblies observed above (56), including the tetramerization property augmented by the G215C mutation.

LLPS of Nref can be induced at higher temperature and by nucleic acid binding (41). It is preceded by the formation of ∼0.1 – 1 μm sized clusters (37, 57). This is accompanied by structural transitions by CD (37), which we similarly observe for N:G215C (SI Appendix Fig. S9). However, N:G215C exhibits much steeper transitions, and at a lower transition temperature, as may be discerned from the temperature-dependent particle size in DLS (Fig. 5i). Interestingly, while for Nref a lower phase transition temperature is achieved in the presence of T_10_, the same transition temperature is observed for N:G215C already without any NA, and addition of T_10_ to N:G215C does not lead to a further shift. Removal of disulfide bonds in reducing conditions further lowered the phase transition temperature. These results indicate more cooperative assembly with lower energy barrier. The largest objects in the co-assembly process of N-protein and NA *in vitro* are droplets from LLPS that are visible in light microscopy. At 20°C (i.e. below the transition temperature) Nref and N:G215C only exhibited small differences, with slightly larger droplets observed for reduced N:G215C compared to Nref (SI Appendix Fig. S10).

### Structural Basis of Protected Islands in Disordered Regions and Effects of the G215C mutation

While several NMR and crystallographic structures are available for the NTD and CTD domains (14, 43, 44, 58–60), the features of the structure and dynamics of the disordered linker and arms are less well understood. The mutationally protected sequence islands in the C-arm at 390–394 and 403–408 and the central linker region 218–231 coincide with transient helices previously revealed in MD simulations (38). As described above, helices in these segments are also found with different structure prediction methods (Fig. 6a and SI Appendix Fig. S3).

**Figure 6.**
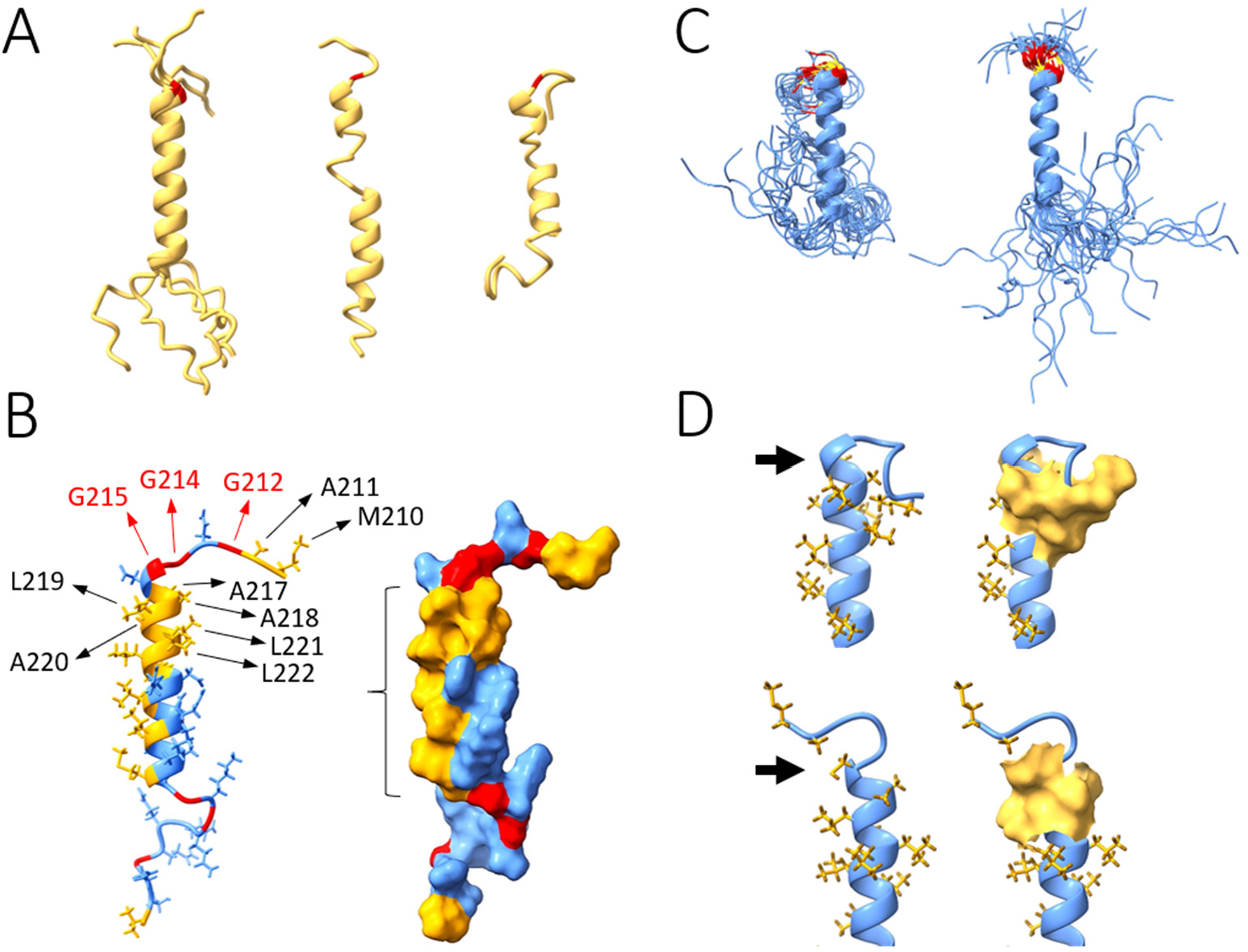
Dynamics simulations of the leucine-rich linker segment 210–246 with and without G215C mutation. **a** Conformations of the reference sequence predicted by AlphaFold2 (left; five models obtained; helix superimposed for comparison), the highest-ranked I-TASSER model (middle), and Phyre2 (right). G215 is highlighted in red. **b** Results of MD simulations for the reference sequence. Left: Positions of residues that play a role in i) stabilizing the helix (mainly by hydrophobic interactions; yellow), ii) conferring flexibility to the N- and C- segments (mainly through the six Gly residues flanking the helix; yellow), and iii) inducing and stabilizing the major changes observed upon the G-to-C mutation. Right: Molecular surface showing the exposed bed of hydrophobic residues (indicated with bracket) likely involved in protein-protein contacts. **c** Snapshots throughout the 100 -nsec dynamics simulation of the C215 (left) and G215 (right) peptides, showing the major conformational changes induced by the mutation. The helix remains structurally stable throughout the simulations regardless of temperature in both cases, but in the mutant, the N-term of the helix is redirected (upward in the figure) and becomes less flexible (position 215 shown in yellow, 214 and 216 in red). **d** Stabilizing interactions. In the reference sequence (upper structures), the flexibility conferred by G215 (arrow) allows M210 and A211 to get close to A218, L219, L221, and L222. These hydrophobic interactions (yellow surfaces) stabilize the N-term segment in a different orientation than in the C215 mutant (lower structures). Here, the Cys sidechain (arrow) shows persistent hydrophobic interactions with A218 and L219, thus redirecting the chain in a different orientation relative to the helix.

To assess the impact of the G215C mutation, we studied the disordered stretch 210–246 in the central linker containing the leucine-rich region. Although structure prediction is generally poor in the disordered segments, all the models show varying degrees of α-helical structure in the sequence of interest, with G215 at the stem of the helix. Highest reliability is obtained in the 222– 234 stretch (Fig. 6a). The top-ranked AlphaFold2 model was used here as the initial structure for MD calculations.

Key residues of the helix can be discerned in Fig. 6b, which highlights six flanking Gly residues conferring structural flexibility, and a bed of hydrophobic residues that stabilize the helix and may serve as a protein-protein interaction interface. Mutation of G215 by C215 results in redirection and reduced flexibility of the N-term of the helix (Fig. 6c). These structural and dynamic changes of the mutant originate both in the higher flexibility of Gly relative to Cys and in the stabilization of the Cys sidechain through persistent hydrophobic interactions with A218 and L219 (Fig. 6d). This results in a more open configuration that appears better poised for helix-helix and thiol interactions. This is consistent with the experimentally observed enhanced dimer-dimer interactions of the N:G215C under reduced conditions and the possibility of forming disulfide bonds across different protomers.

## Discussion

The worldwide sequencing effort has led to the assembly of an unprecedented database of viable SARS-CoV-2 variants, alone for N-protein describing ∼25,000 different species as of November 2021. In the present work we have combined an analysis of the amino acid variability of SARS-CoV-2 N-protein with biophysical experiments of select mutants, and found remarkable plasticity on all levels of organization. Plasticity has been hypothesized to be a unique feature of RNA virus proteins, which have more loosely packed cores and an abundance of intrinsically disordered regions that confer high degrees of flexibility for adaptation and tolerance to mutations (25). In the present case of the SARS-CoV-2 N-protein, more than 86% of positions in the amino acid sequence are subject to variation, on average by 3-4 different amino acids, about half of which score low on a physicochemical similarity scale. We observed substitutions at many positions thought to be critical due to their conservation across related coronaviruses, or their constitution of phosphorylation and protein interaction sites, are found to be viable. Our data show that the single mutation G215C causes significantly altered secondary structure; gross differences in hydrodynamic shape indicate altered subunit arrangements; and strongly enhanced protein-protein interactions modulate the formation of quaternary structure. More extended studies are required to elucidate the expected impact of mutations in N-protein, for example, on host protein interactions, interactions with other viral proteins, and on phase boundaries for condensation and nucleic acid coassembly. This plasticity poses significant challenges to identify the essential functions and mechanisms that may serve as therapeutic targets in N-protein.

Earlier examinations of emerging mutations in N proteins (6, 14, 61–63), going back to June 2020, were necessarily more limited in scope, and while sufficient to examine hot spots and identify key replacements such as R203K/G204R (6, 10, 14, 64), it was not yet possible to draw conclusions from a survey of the entire mutational landscape. Due to the orders of magnitude larger coverage that has become available through the global genomic epidemiology efforts since then, we believe that the observed data now approaches the limits of possible mutations for functioning N-protein, and therefore reflects its biophysical properties.

The study of the constraints in the mutational landscape appears very useful to complement structural biophysical tools, particularly to examine intrinsically disordered regions. These generally are permissive for a wide range of amino acid substitutions and, in fact, harbor three of the four persistent N-protein mutations characteristic of the Delta variant, and all of the Omicron mutations in N-protein. However, the mutational landscape of N-protein reveals several islands within these regions that are highly protected and therefore appear to have critical functions.

One of these is in the central linker adjacent to the G215 position, where the G215C mutation has quickly outcompeted all other variants in 2021 after its appearance alongside only ORF1ab and ORF7b mutations in the Delta variant. NMR and molecular dynamics studies have reported distinct α-helical propensity flanking position 215 in the SR-rich region (63) and in the leucine-rich region (38). The latter was proposed by Cubuk et al. to provide interfaces for oligomerization. Most recently, NMR experiments showed formation of an α-helix 219-230 when in complex with nsp3a, which binds to N-protein competitively with RNA (65). In the present work, we found this region highly protected against mutation. We propose that these helices may be essential for higher-order assembly of N-protein and are either stabilized or exposed in the emerging G215C mutant. Indeed, our simulations show a stable helix spanning residues G215-S235 exhibiting an extended hydrophobic surface on one side. Mutation to C215 repositions the downstream sequence relative to the helix, which renders the hydrophobic surface more accessible for protein-protein interactions.

The potential introduction of disulfide bonds in the N:G215C mutant stabilizing dimer-dimer crosslinks in the linker region would constitute an even more drastic change in the assembly pathway. It is uncertain, however, whether these disulfide bonds are formed *in vivo*. Disulfide bonds are absent in nucleocapsid proteins of related betacoronaviruses, but assist oligomerization of nucleocapsid proteins of other RNA viruses at least transiently (66, 67). However, even without disulfide bonds, we found a 200-fold enhancement of dimer-dimer self-association in N:G215C, accompanied by augmented co-assembly with NA, which we hypothesize profoundly alters the co-assembly kinetics of ribonucleoprotein particles. This may contribute to the clinical phenotype of strongly increased viral load and infectivity of the Delta variant, and the overwhelming dominance of the 21J Delta clade relative to the 21I Delta clade despite the lack of additional changes in the spike protein.

After writing of this manuscript, the 21J Delta variant of SARS-CoV-2 has been replaced by the Omicron variant as the worldwide dominant variant. Interestingly, the latter has none of the N-protein mutations defining for the Delta variant, but it exhibits deletions and new mutations that may impact viral fitness, in addition to the characteristic spike protein mutations. While this development does not impact our conclusions on the mutational landscape of N-protein and its relation to its biophysical properties and assembly functions, it will be interesting to examine to which extent Omicron will explore, on a global population level, a similar N-protein sequence space as Delta and prior variants.

## Materials and Methods

### Sequence analysis

Mutation data were based on sequenced SARS-CoV-2 genomes submitted to the GISAID (Global Initiative on Sharing All Influenza Data), and downloaded on November 29, 2021 as preprocessed file by the Nextstrain team (nextstrain.org) (13) containing 2.49 million sequences. 49.7% of all sequences included the N:G215C mutation characteristic for the dominant Delta clade, using the Wuhan-Hu-1 isolate (GenBank QHD43423) (68) as the ancestral reference. To define a threshold for sequence errors, the daily mutation events in positions that are strictly conserved across coronaviruses were inspected. While 7 exhibited no mutations at all since January 2020, and 6 only a single event, those that were mutated in excess of 10 times exhibited clusters of two or more closely spaced events as would be expected from transmission chains. For positions with more than 20 total mutation events, clusters occurred frequently to approach a virtually continuous accumulation of their total count. Thus, a threshold of 10 observations was set as a lower limit for any mutations to be considered in the present work.

Alignment of SARS and related sequences was carried out with COBALT at NLM (69), and highlights for similar residues were taken from ESPript (70), plotted with MATLAB (Natick, MA). Similarity scores were calculated using the EMPAR matrix (71). Propensity of residues to promote LLPS were calculated by FuzDrop (72).

### Molecular modeling and dynamics simulations

Three dimensional models of the sequence ^210^MAGNGGDAALALLLLDRLNQLESKMSGKGQQQQGQTV^246^ were obtained from three independent servers that use different assumptions and algorithms (73–75). The model used here was obtained with AlphaFold2 after extracting the predicted coordinates of the above sequence from the modeled full-length N-protein. All the models predict modest helical content, except for the N- and C-term segments, which are disordered. Structures of the C-term helix were also obtained with the three prediction methods based on the sequence ^390^QTVTLLPAADLDDFSKQLQQSMSSADSTQA^419^.

MD simulations were carried out in the NPT ensemble, at 25 °C and 37 °C and 1 atm, in a cubic cell with PBC and PME summations, using the all-atom CHARMM (param36) force field (76, 77). The peptides were capped (acetylated N- and amidated C-) to minimize potential electrostatic artifacts of the termini. All bond lengths involving hydrogen atoms were constrained with the SHAKE algorithm, and an integration step of 2 fs was used. The pressure was maintained with the Langevin piston method, with mass and collision frequency of 400 amu and 20 ps^−1^. The temperature was maintained with the Hoover thermostat, using a mass of 10^3^ kcal mol^−1^ps^2^. The side length of the simulation box was initially set at ∼9.3 nm and filled with ∼27,000 TIP3P water molecules, yielding an average density of ∼0.993 g/cm^3^ at 37 °C after equilibration. Assuming Asp^−^ and Glu^−^ unprotonated and Arg^+^ and Lys^+^ protonated at neutral pH, the peptides are electroneutral; 74 K^+^ and 74 Cl^−^ ions were added to mimic near-physiological [KCl] ∼150 mM concentration. The ions were randomly distributed in the water phase after the peptides were solvated and the overlapping water molecules removed. After standard protocols of heating and equilibration, a productive phase of 100 ns was conducted, and analysis performed over the last 80 ns. Structural and dynamic analyses were based on the calculations of average values and standard deviations of the ϕ and ψ dihedral angles per residue.

### Protein and oligonucleotides

SARS-CoV-2 nucleocapsid protein # YP_009724397 with quadruple D63G, R203M, G215C, and D377Y mutations including 6His with TEV cleavage site was synthesized and cloned into the pET-29a(+) expression vector by GenScript (Pisctaway, NJ). The plasmid was transformed into BL21(DE3)pLysS *E*. *coli* (ThermoFisher catalog # C606010), and grown in LB kanamycin at 37°C to 0.8 OD. Protein expression was induced with 0.5 mM IPTG overnight at 18°C. Cells were harvested and then lysed in 20 mM Na_2_PO_4_ pH 7.5, 1.5 M NaCl with 1 tab of protease inhibitor cocktail (SIGMAFAST, Sigma-Aldrich, St. Louis, MO; catalog #S8830) passing twice through an Emulsiflex-C5 extruder (Avestin Inc., Ottawa, ON, Canada) followed centrifugation at 4°C to remove cell debris. A Ni^2+^ affinity column (HisTrap FF Crude 5 ml, Cytiva, Marlborough, MA) was equilibrated in 20 mM Na_2_PO_4_ pH 7.5, 1.5 M NaCl and cell lysate was added at a flow rate of 0.5 mL/min. The column was washed in the same buffer supplemented with 20 mM imidazole. Following a modified unfolding/refolding protocol by (40) to remove residual protein-bound bacterial nucleic acid, the captured protein was denatured in 50 mM HEPES pH 7.5, 500 mM NaCl, 6 M urea, 10% glycerol for at least 20 column volumes at 1 mL/min, and then renatured in an overnight gradient wash with 40 column volumes at a flow rate of 0.2 mL/min to a final buffer 50 mM HEPES pH 7.5, 500 mM NaCl, 10% glycerol. Protein was eluted in a gradient to 50 mM HEPES pH 7.5, 500 mM NaCl, 10% glycerol, 500 mM imidazole at 0.2 mL/min. For cleavage of the 6His tag, peak fractions were quickly dialyzed against cleavage buffer of 20 mM Na_2_PO_4_, 250 mM NaCl, 1 mM DTT, pH 7.5 to remove imidazole, and then incubated overnight with TEV protease in cleavage buffer supplemented with 0.5 mM EDTA. Reaction products were dialyzed in 50 mM HEPES, 500 mM NaCl, pH 7.5 and then purified by elution through Ni^2+^ affinity column. Protein concentration was measured by UV/VIS spectrophotometry, and an absorbance ratio at 260 nm to 280 nm of ∼0.55 was observed, confirming the absence of nucleic acid. Protein purity was confirmed by SDS-PAGE.

SARS-CoV-2 N-protein accession # YP_009724397, the N:G215C mutant and the N:D63G mutant were acquired from EXONBIO (San Diego, CA; catalog# 19CoV-N150, 19Cov-N180, and 19Cov-N170). Both constructs have a C-terminal His-tag and were expressed in *E*. *coli*. Sequences and the absence of post-translational modifications were verified by LC-MS/MS. For Nref, based on amino acid composition the molar mass is 46,981.06 Da, and the molar extinction coefficient at 280 nm is 43,890 M^−1^cm^−1^. For N:G215C the MW is 47027.15 Da and for N:D63G it is 46,923 Da . All have a predicted partial-specific volume of 0.717 ml/g at 20°C. The protein was formulated in phosphate buffer pH 7.4, 250 mM NaCl and stored in frozen form. The ratio of absorbance at 260 nm to 280 nm of Nref, N:D63G and N:G215C was 0.52, 0.56 and 0.56 respectively. Prior to biophysical characterization the proteins were dialyzed exhaustively the working buffer (Na_2_PO_4_ 10.1 mM, KH_2_PO_4_ 1.8 mM, KCl 2.7 mM, NaCl 10 mM, pH 7.40). Final protein concentrations were determined by spectrophotometry or refractometry.

The oligonucleotides T_6_ (TTTTTT), T_10_ (TTTTTTTTTT) and T_20_ (TTTTTTTTTT TTTTTTTTTT) were purchased from Integrated DNA Technologies (Skokie, IL), purified by HPLC and lyophilized. They were dialyzed in the working buffer and their concentration measured by absorption spectrophotometry as previously described (37).

### Analytical ultracentrifugation

Sedimentation velocity (SV) experiments were carried out in a ProteomeLab XL-I analytical ultracentrifuge (Beckman Coulter, Indianapolis, IN) as previously described (78). Protein samples were loaded in cell assemblies comprising charcoal-filled Epon double-sector centerpieces of 3 mm or 12 mm pathlength and sapphire windows. The samples were temperature equilibrated at 20°C in an AN-50 TI rotor, followed by acceleration to 50,000 rpm. Depending on the solution composition of the samples, data were acquired with Rayleigh interference optics and absorbance optics at 230 nm, 260 nm, and/or 280 nm. Standard protocols were followed to calculate the sedimentation coefficient distribution *c*(*s*) for each data set using the software SEDFIT (79). For the concentration series of N:G215C, the integrated weight-average sedimentation coefficients, *s*_w_ were assembled into *s*_w_ isotherms and modeled with a monomer-dimer equilibrium in the software SEDPHAT (53).

### Dynamic light scattering

Autocorrelation data were collected in a NanoStar instrument (Wyatt Technology, Santa Barbara, CA). 100 μL samples at 3 μM N-protein in the presence or absence of oligonucleotides were inserted into a 1 μL quartz cuvette (WNQC01-00, Wyatt Instruments), using excess sample to minimize impact of evaporation in the observation chamber. Laser light scattering was measured at 658 nm at a detection angle of 90°. For the temperature scans, a ramp rate of 1°/min was applied with 5 sec data acquisitions and averaging 3 replicates for each temperature point. Data were collected and processed by using software Dynamics 7.4 (Wyatt Instruments) or SEDFIT (National Institutes of Health).

### Mass photometry

The mass distribution of N:G215C was determined using a Refeyn One instrument (Refeyn, Oxford, UK). The measurements followed the standard protocol (80). Briefly, 10 μL of freshly filtered buffer was loaded in a well of a gasket (CultureWell, GBL103250, Sigma, MO, USA) for image focusing, and then 10 μL of protein solution was added to a final concentration of 10 nM and mixed by pipetting. Immediately after the mixing, a 1-minute video was recorded using the AcquireMP software (Refeyn, UK). The video was processed by using the DiscoverMP software (Refeyn, UK) and the contrast value of each protein molecule was converted to mass with calibration obtained from an unstained protein ladder (LC0725, Thermofisher, Wattham, MA).

### Circular dichroism spectroscopy

CD spectra were acquired in a Chirascan Q100 (Applied Photophysics, U.K.). Samples were measured in 1 mm pathlength cells, with 1 nm steps and 1 sec integration time. Results are averages of 3 acquisitions. Backgrounds of corresponding buffers (with or without oligonucleotides, respectively) were subtracted. For temperature scans, data were acquired in 1 nm intervals with integration times of 0.5 sec, without repeats, applying a temperature ramp rate of 1°C/min. The Global3 software from Applied Photophysics was used to deconvoluted the multi-wavelengths temperature scans.

### Microscopy of *in vitro* liquid-liquid phase separated condensates

The *in vitro* phase separation assays were performed at 23 °C. N protein was studied in the presence of oligonucleotides T_10_ or T_20_ at different concentrations in the working buffer (Na_2_PO_4_ 10.1 mM, KH_2_PO_4_ 1.8 mM, KCl 2.7 mM, NaCl 10 mM, pH 7.40) or working buffer supplemented with 1 mM TCEP. Samples were mixed in 1.7 mL microcentrifuge tubes and then immediately transferred onto a glass-bottom 35 mm dish (catalog # Part No: P35G-1.5-20-C, MatTek). Condensates from LLPS were imaged within 30 – 40 min. Images were acquired on a Nikon Ti-E microscope equipped with a Prime 95B camera (Teledyne Photometrics) with sensor dimensions of 1200×1200 pixels. Images were collected using a 100X 1.49 NA oil objective lens with a pixel size of 110 nanometers. The transmitted light source was a collimated white light LED (Lumencor PEKA) passed through a green interference filter.

### Differential scanning fluorometry

Thermal scans with measurement of the intrinsic fluorescence of the protein samples were carried out using a Tycho instrument (Nanotemper, Germany). 10 μL samples were loaded in capillaries (TY-C001, Nanotemper). The intrinsic protein fluorescence was measured at 350 nm and 330 nm, and the first derivative of the intensity ratio was calculated as a function of temperature. The temperature ramp rate was 30°C/min and data were acquired from 35 to 95 °C.

## Acknowledgments

We thank Drs. Dominic Esposito, Myungwoon Lee, Robert Tycko, and Harshad Vishwasrao for helpful discussions and sharing of resources. This work was supported by the Intramural Research Programs of the National Institute of Biomedical Imaging and Bioengineering, the National Institute of Allergy and Infectious Diseases, the National Heart, Lung, and Blood Institute, and the National Institute of Neurological Disorders and Stroke, National Institutes of Health. This work utilized the computational resources of the NIH HPC Biowulf cluster for sequence analyses and dynamics simulations.

## Supplementary Information for

### Supplementary Information contains

**Tables S1 and S2**

**Figures S1 to S10**

**Table S1.** Scope and depth of mutations in SARS-CoV-2 proteins. Sequence data from GISAID, pre-processed and downloaded from Nextstrain.org on November 29^th^ 2021, was parsed for the occurrence of mutations at each protein. Mutations were counted that appeared above a threshold of 10 times. At each position, the number of distinct substitutions was recorded and averaged over all positions that show any substitutions.

| **Protein** | **% of positions with mutations** | **average # of substitutions** |
| --- | --- | --- |
| E | 77.3 | 2.48 |
| M | 71.6 | 2.28 |
| N | 86.4 | 3.49 |
| S | 79.3 | 2.87 |
| ORF1a | 86.9 | 2.67 |
| ORF1b | 75.6 | 2.33 |
| ORF3a | 97.8 | 4.02 |
| ORF6 | 100 | 3.60 |
| ORF7a | 100 | 4.43 |
| ORF7b | 100 | 3.66 |
| ORF8 | 98.4 | 3.74 |

**Table S2.** Serine mutations in the SR-rich sequence. The number of serines occurring at positions 176-206 is counted in each of the 24,957 unique N sequences. In the ancestral sequence the number is 14. We count as a deletion of serine the mutation of a serine at specific position to any other amino acid. Of all sequences, 14.3% have at least one serine deleted, but of these only 2.28% have a second deletion. Thus, probabilities of deletion of serine are significantly lower for sequences with existing deletions (χ^2^ = 315 (f=1), P < 1e-8), showing anti-correlation of serine mutations. *Vice versa*, insertion of a serine at a new position occurs only 2.42% of the time overall.

| sequence | # of serine residues | # of sequences | % of sequences |
| --- | --- | --- | --- |
| no change (ancestral) | 14 | 21,388 | 85.7% of all |
| any change |  | 3,569 | 14.3% of all |
| ≥1 S deletion | ≤13 | 3027 | 12.1% of all |
| 2 S deletions | 12 | 69 | **0.3% of all** |
|  |  |  | **2.3% of ≥1 deletion** |
| > 2 S deletions | <12 | 0 | 0% |
| ≥1 new S position | ≥15 | 603 | 2.4% of all |
| 1 S deletion + ≥1 new S | ≥14 | 61 | **0.24% of all** |
|  |  |  | **2.0% of ≥1 deletion** |

**Figure S1.**
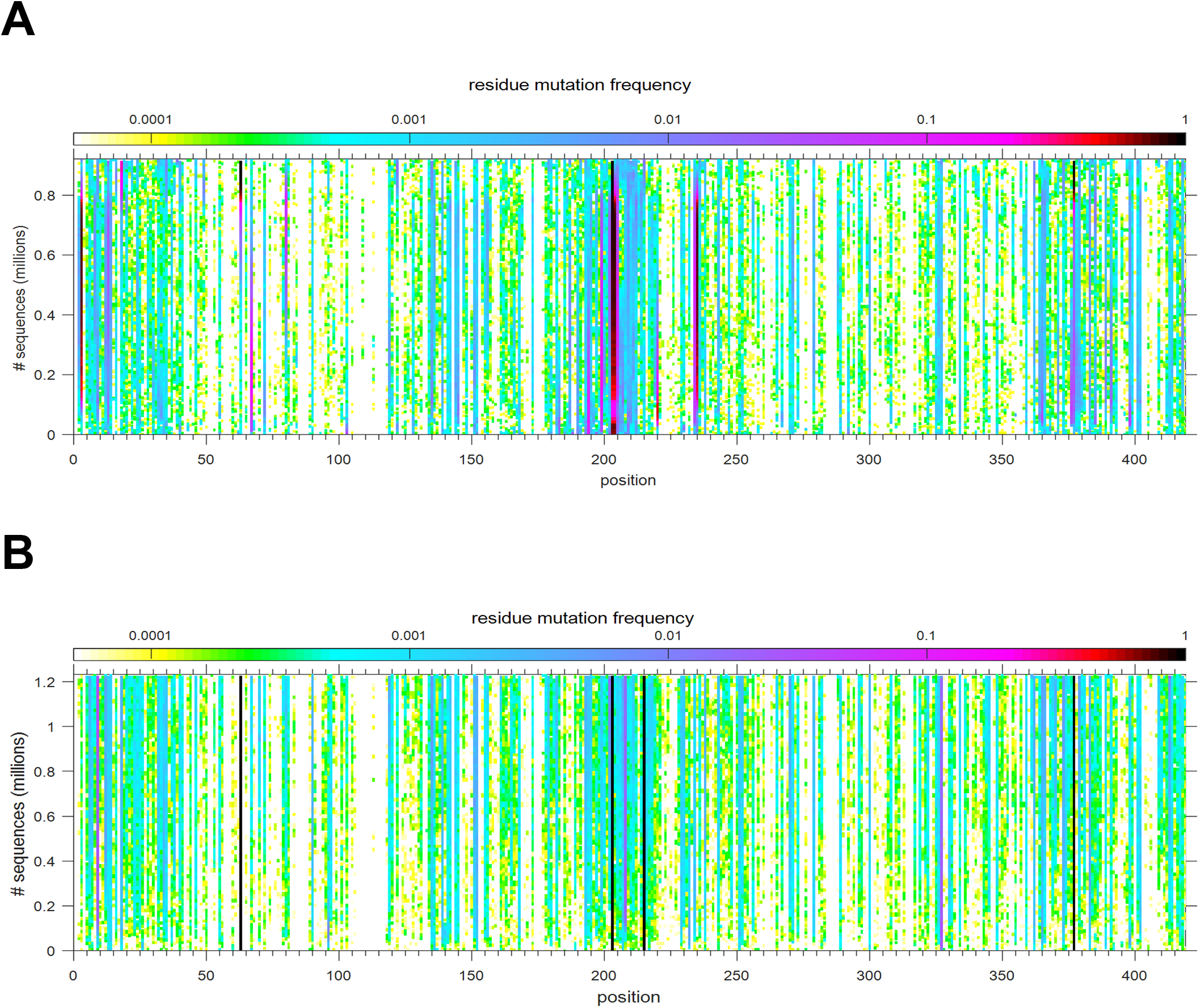
Temporal rate of mutations of variants. In an analogous presentation to **Fig. 3e**, shown is the frequency of mutations *vs.* total sequence number, subdivided for variants preceding Delta 21J (**A**) and Delta 21J (**B**). For each position, the daily number of any mutations relative to the number of new sequences, convoluted across 10^4^ sequences, is plotted against the total accumulated sequences (as scale of time) and color-coded according to the relative rate of observing a mutation. The initial spread in the mutational landscape in shown **Fig. S2**.

**Figure S2.**
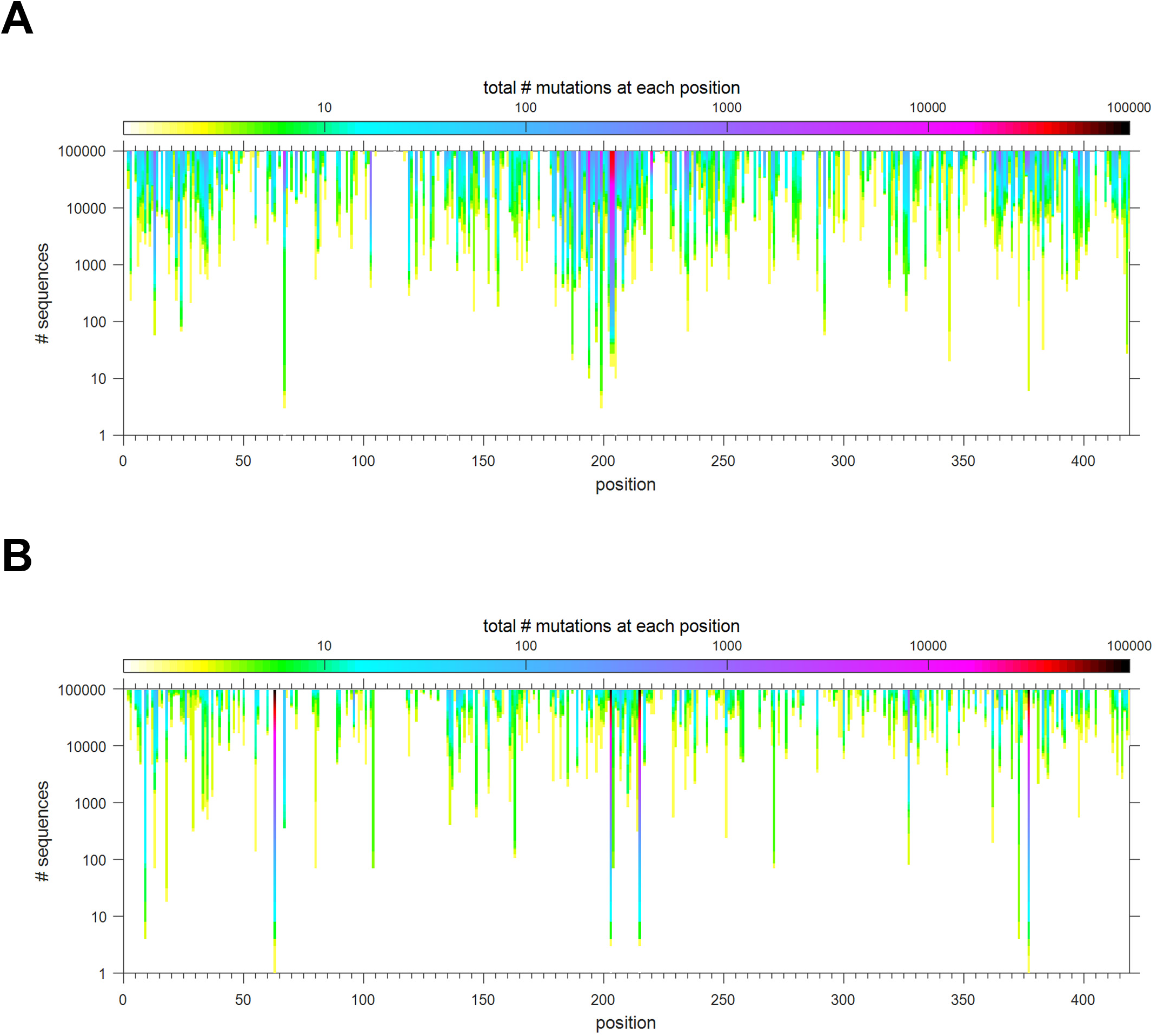
Spread of observed instances of mutations across sequence space. Shown is the accumulation of observed instances of mutations in each position, as a function of total number of sequenced genomes, for the first sequences of SARS-CoV-2 (**A**), and for Delta 21J (**B**). Initially only the most frequent mutations have been recorded, and more rare mutations are observed after many genomes have been sequenced.

**Figure S3.**
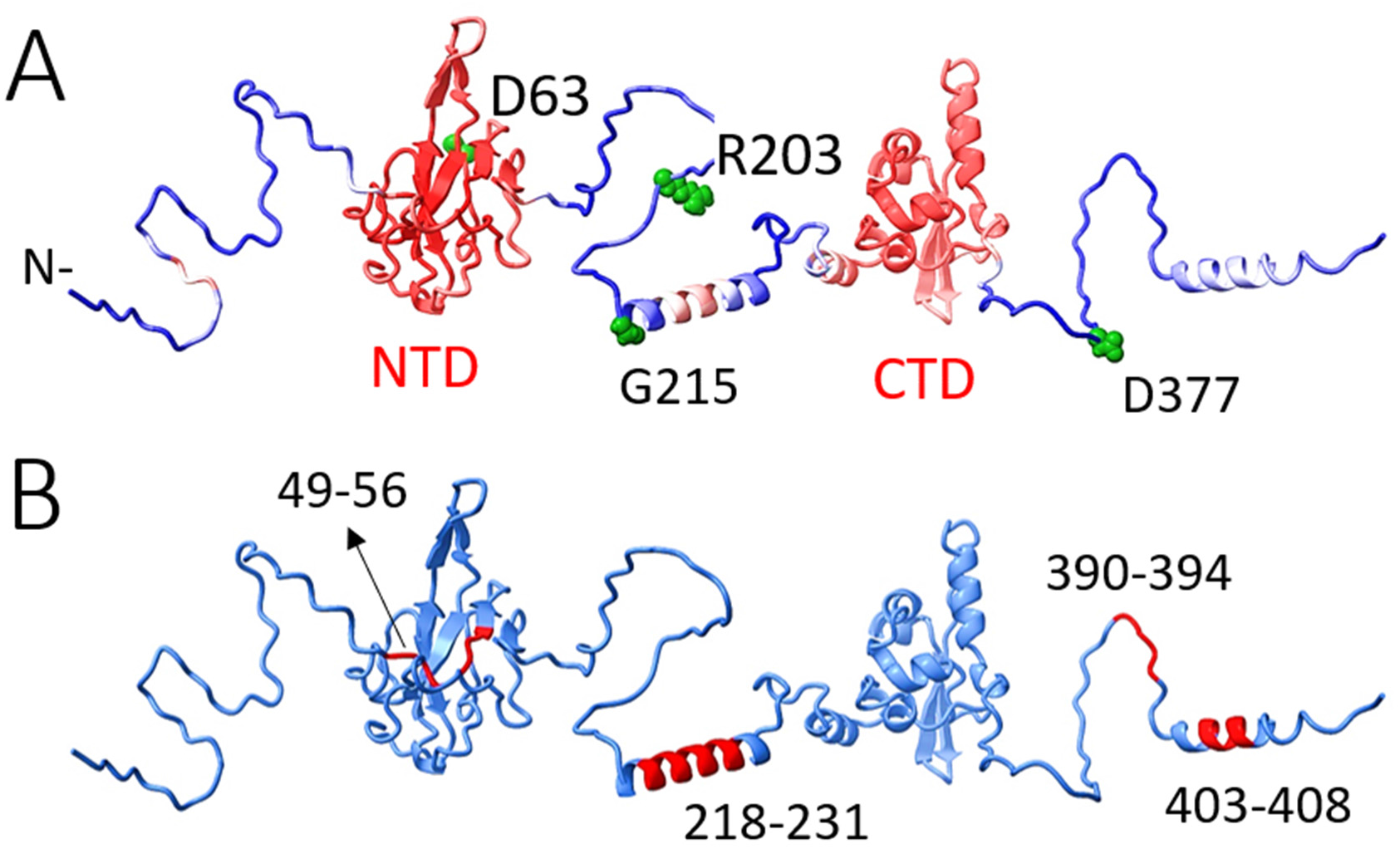
Three-dimensional model of full-length N-protein. (**A**) AlphaFold2 model of the N-protein (ancestral sequence) color-coded according to the prediction reliability (red: highest confidence; blue: lowest). While CTD and NTD are mostly red, the disordered regions (N-arm, linker, and C-arm) are mostly blue but show two short segments with helical content: G215-S235 and L400-S410. Similar helices in both positions are produced by independent calculations in structure prediction servers I-TASSER and Phyre2, albeit with low confidence, suggesting an intrinsic propensity for these short sequences to form helices within the highly disordered segment. Critical residues observed in the Delta variant are highlighted in green. The backbone of the disordered segments was rearranged relative to the folded domains for visualization purpose. (**B**) AF2 model showing in red the four regions most protected from mutations (yellow patches in Figure 3).

**Figure S4.**
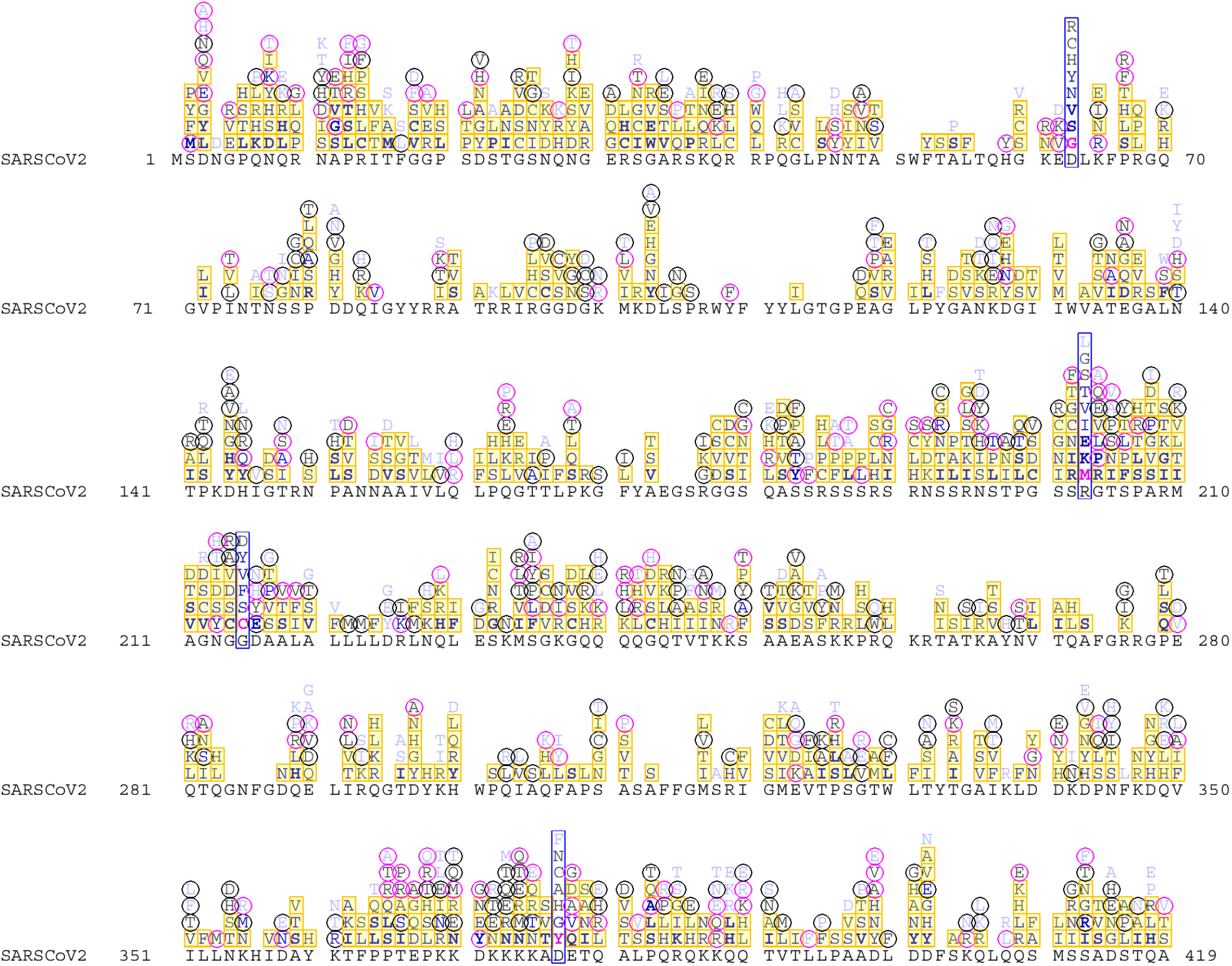
Amino acid substitutions in the mutational landscape of 21J Delta SARS-CoV-2 and non-21J Delta variants. The ancestral reference sequence Wuhan-Hu-1 is depicted in black letters, above which the range of observed mutations is presented, as in Extended Data Fig. 1 with different weights for different number of observations 10–20 (light blue), 20–100 (gray), 100-1,000 (blue), and > 1,000 times (bold blue). Mutations that occur in both 21J and non-21J sequences are highlighted in yellow squares, those occurring only in sequences from the 21J clade are encircled in black, and those only observed in non-21J sequences are encircled in magenta. The blue bars highlight the position of the four mutations characteristic for 21J Delta sequences. Mutations that have no highlight indicate rare mutations that only jointly exceed the threshold of 10 reports, but not in either the 21J or non-21J group alone.

**Figure S5.**
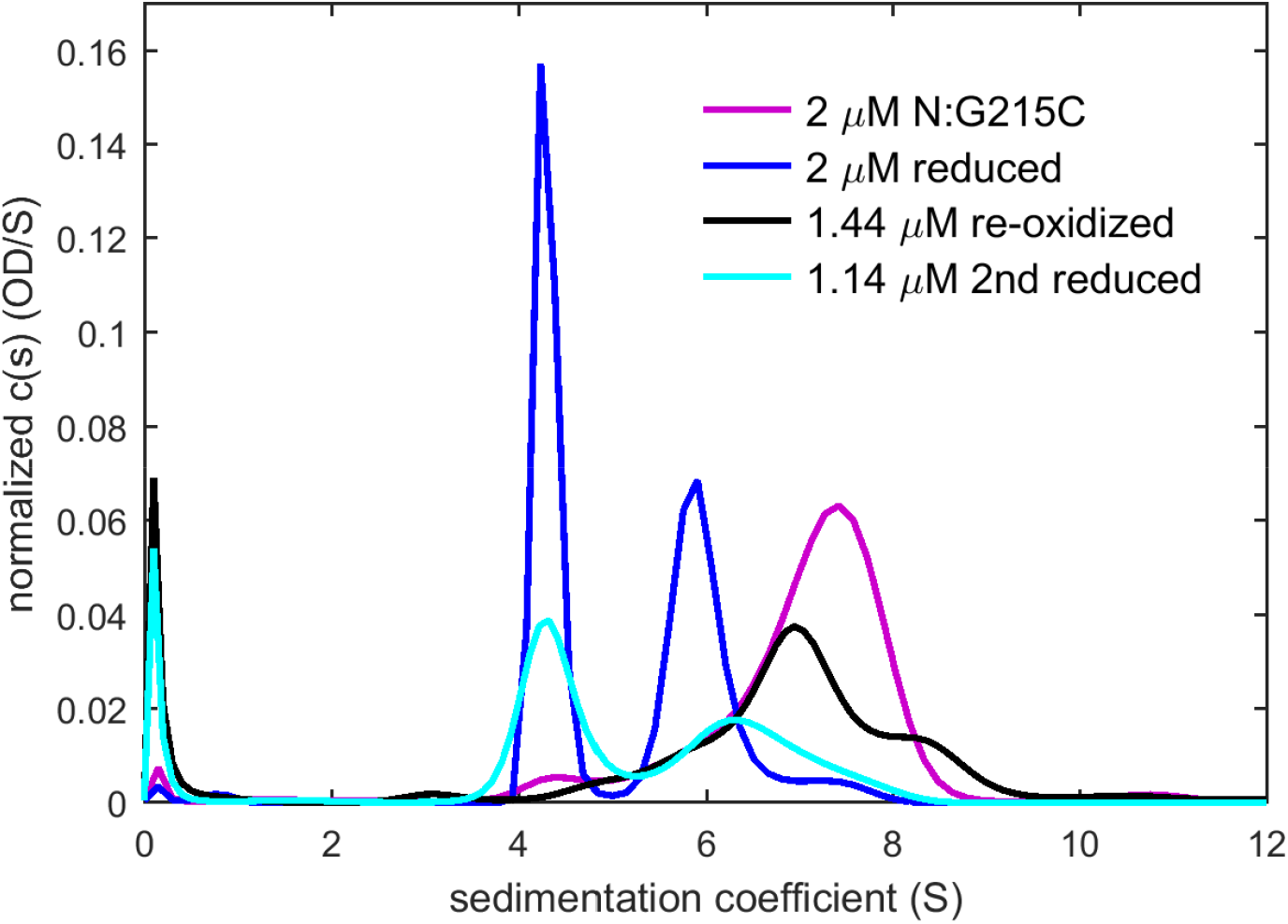
Oligomeric state of N:G215C after reversible reduction and re-oxidation. We asked whether the oligomeric distribution under non-reducing conditions might just be dynamically locked into the observed tetramer, and whether it is possible to alter it, if disulfide bonds are broken and then reformed. Initially, the original diluted protein stock is in a disulfide-linked tetrameric state (magenta), in a replicate experiment of **Fig. 5a**. Next, after dilution into 1 mM TCEP and incubation for 2 hours (blue) the majority of the protein is in a dimeric state (linked by CTD interactions) and exhibiting a reversible dimer-tetramer self-association equilibrium, replicating results of Fig. 3b. This is followed by removal of TCEP by overnight dialysis during which the protein is allowed to re-oxidize (black), which produces an assembled state, with an average s-value of 7.03 S virtually identical to the original state with 7.07 S. Based on hydrodynamic scaling laws, a hexamer would be expected to sediment at ∼9.6 S, and a octamer at ∼11.6 S. Importantly, the reformed disulfide bonds did not cause significant populations of oligomers larger than the tetramer. Finally, in order to demonstrate that these re-associated oligomers are disulfide-linked (as opposed to degraded misfolded oligomers), this sample was diluted again into 1 mM TCEP and incubated for 2 hours (cyan), reproducing the majority dimeric reduced protein state.

**Figure S6.**
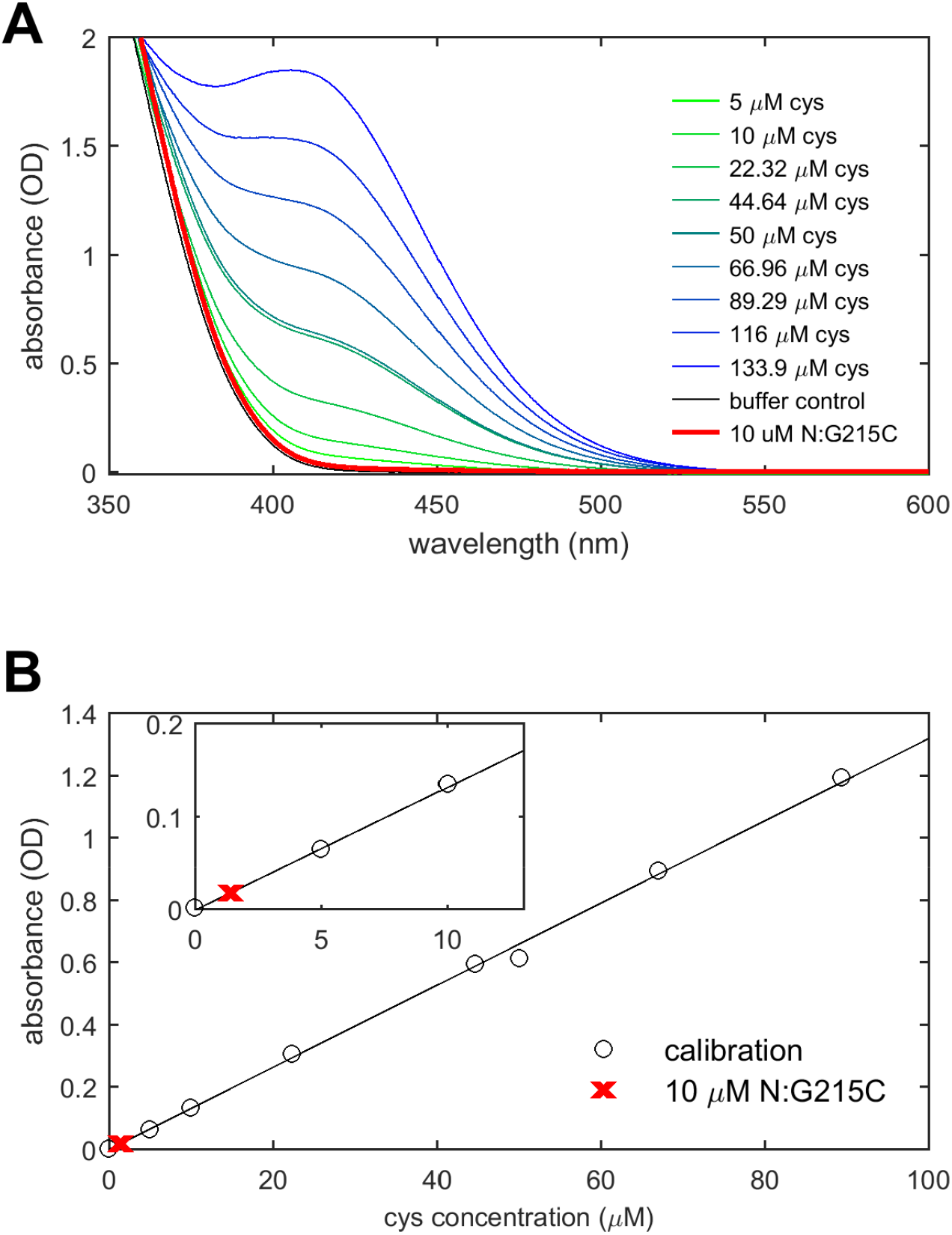
Free thiols assay determining the completion of disulfide bonds in N:G215C. (**A**) Absorption spectra of a concentration series of cysteine solutions after reaction with sulfhydryl reagent 5,5′-dithiobis (2-nitrobenzoic acid) (DTNB) (green to blue); and spectra of a buffer control and 10 μM N:G215C. (**B**) Conversion of spectral maximum at 412 nm into a standard curve (circles) and experimental value for 10 μM N:G215C, leading to an estimate of 1.4±0.1 μM free thiols in 10 μM N:G215C.

**Figure S7.**
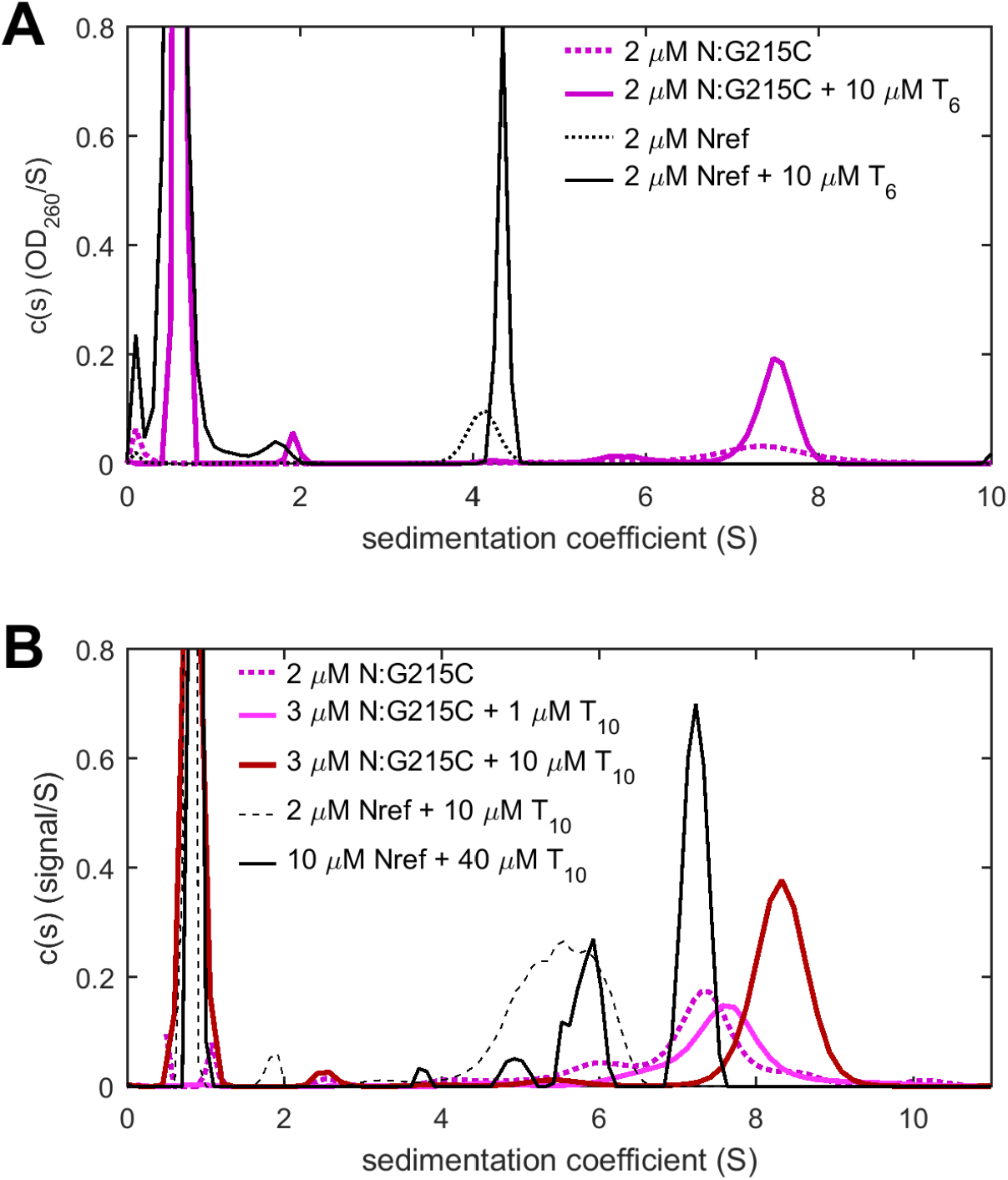
Nucleic acid binding to N:G215C oligomers. Binding is studied by SV using multiple detection systems simultaneously to measure NA binding while monitoring protein oligomeric states. (**A**) For reference, the hexa-oligonucleotide T_6_ can ligand Nref dimers with a K_D_ of 0.6 μM at two sites (black solid and dotted lines for Nref with and without T_6_) (Zhao et al., 2021a). The short oligonucleotide adds only little to the protein mass in the complex, and therefore does not significantly increase the sedimentation velocity. However, binding can be assessed from the increase of the 260 nm absorbance co-sedimenting with the complex. Therefore, sedimentation coefficient distributions c(s) were calculated based on sedimentation profiles acquired at 260 nm. For Nref the 260 nm absorbance increases 2.8fold due to bound T6 at the conditions shown (black solid line). For N:G215C, the increase in 260 nm absorbance is 2.3fold (magenta line), suggesting similar but slightly weaker binding. (**B**) Binding of the oligonucleotide T_10_ to Nref is close to stoichiometric at micromolar concentrations, and induces reversible N-protein dimer-dimer self-association (black lines). For Nref, the sedimentation coefficient distribution reflects time-average populations of dimer and tetramer species in rapid exchange, which exhibits a characteristic concentration dependence (Schuck and Zhao, 2017). For N:G215C, already at low concentrations of 2 μM N:G215C with 1 μM T_10_ (magenta) no free oligonucleotide is left, indicating high-affinity binding of T_10_ to N:G215C similar as to Nref. The oligomeric state of N:G215C liganded with T_10_ is tetrameric, similar to N:G215C in the absence of NA, with a slight shift consistent with the added mass from multiple T_10_ molecules. When applying a large molar excess of T_10_ (red), a larger shift is observed, consistent with added mass from up to eight copies of T_10_ expected to bind to the tetramer.

**Figure S8.**
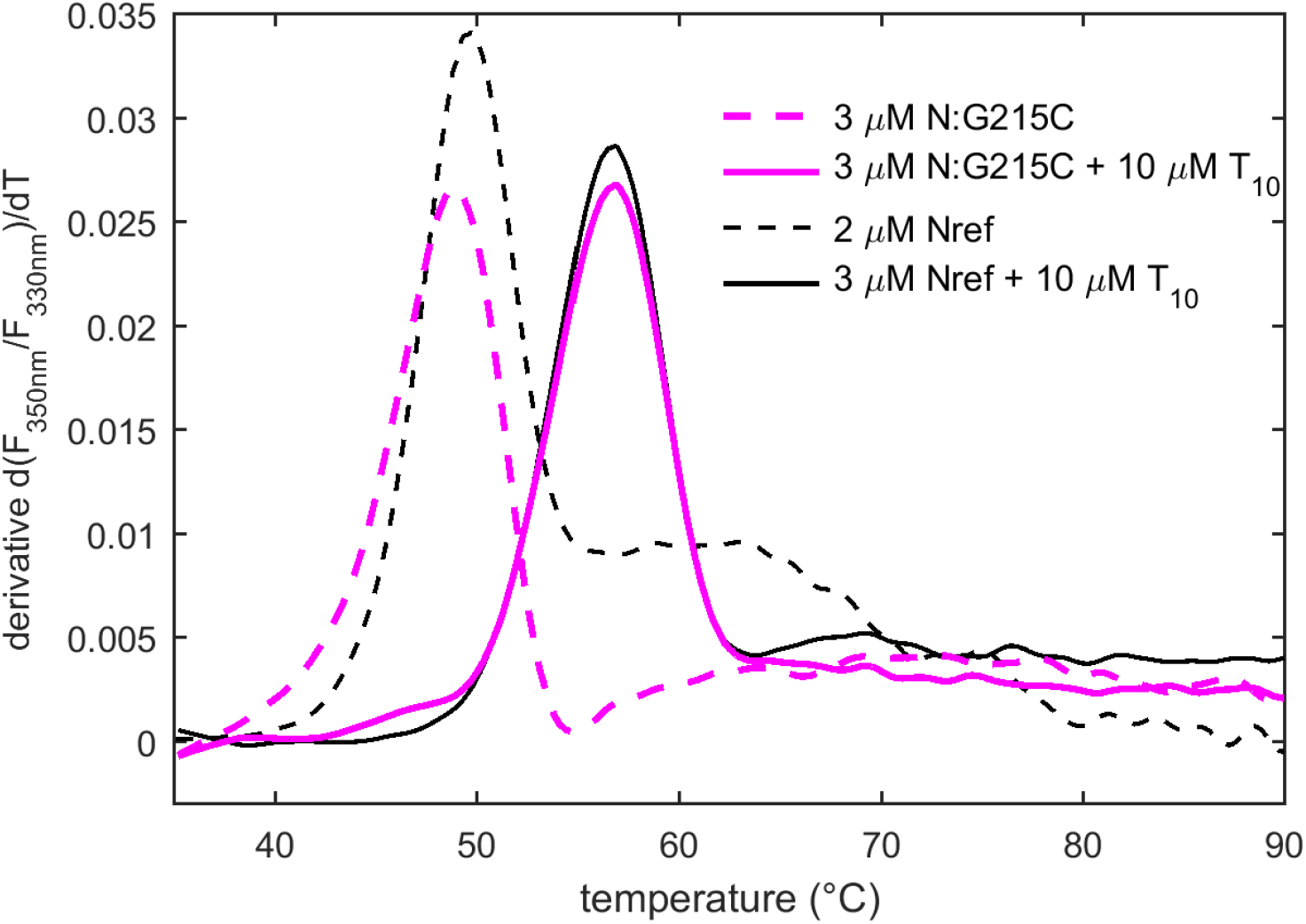
Thermal stability of N-protein and in complex with oligonucleotide T_10_. Shown is the temperature-derivative of the intrinsic fluorescence ratio at 350 nm–330 nm (DSF) for Nref (black) and N:G215C (magenta) with and without molar excess of T_10_. This data reflects on the thermal stability of the folded domains and their immediate vicinity, since tyrosine and tryptophan residues which are contributing to the recorded signal reside exclusively in the folded CTD and NTD domains. The shift to higher transition temperatures in the presence of T_10_ demonstrates higher thermodynamic stability of the complex.

**Figure S9.**
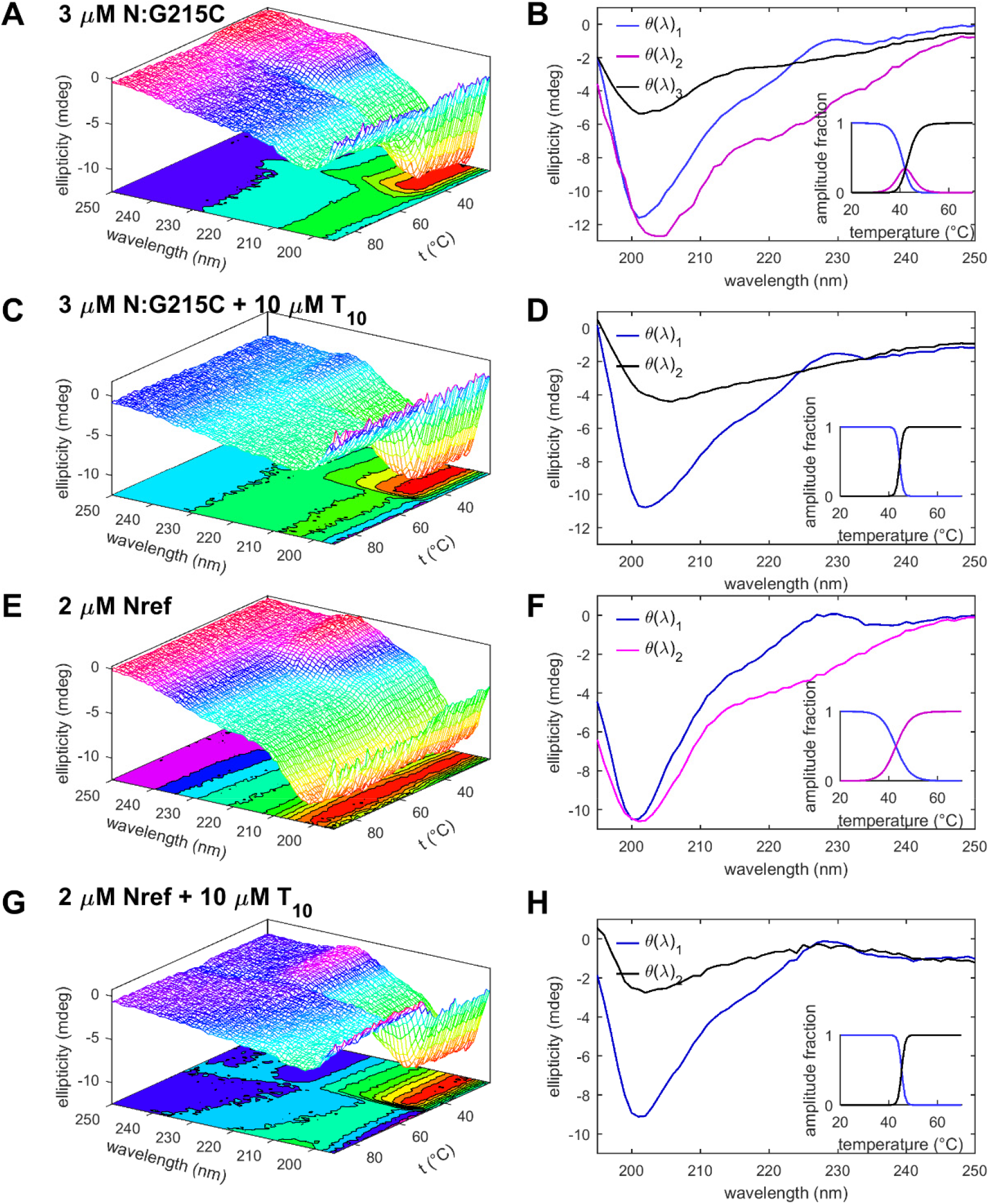
Temperature dependent secondary structure by CD. CD spectra were recorded as a function of temperature (left column) and decomposed into basis spectra (right column) and their relative contributions as a function of temperatures (insets). (**A**, **B**), CD data for 3 μM N:G215C, (**C**, **D**), 3 μM N:G215C with 10 μM oligonucleotide T_10_, (**E**, **F**), 3 μM Nref, (**G**, **H**) 3 μM Nref with 10 μM T_10_. N:G215C in working buffer (**A**, **B**) shows a transition at ∼43°C from a largely disordered state (blue) to a state with increased negative 220-230 nm ellipticity characteristic for helical content (magenta). This mirrors the conformational changes for Nref (**E**, **F**), but for N:G215C it is followed quickly by a loss of signal (black) caused by depletion of material from the light path due to sedimentation of large particles formed. Similar depletion is observed for all CD data in the presence of T_10_. Data for Nref are reproduced from Zhao et al. (Zhao et al., 2021a).

**Figure S10.**
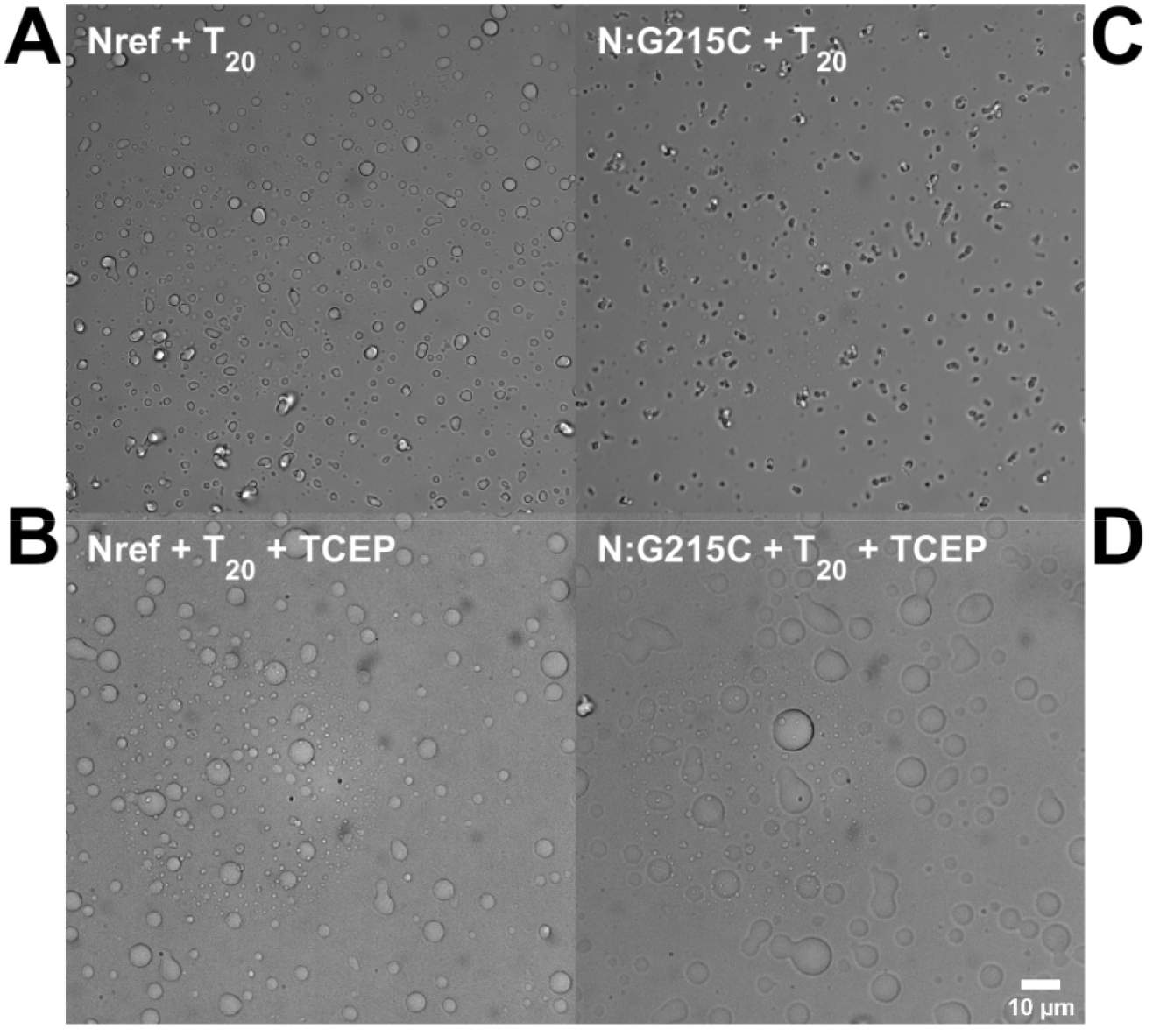
LLPS and particle formation of SARS-CoV-2. Shown are brightfield microscopy images of 5 μM Nref (left) and N:G215C (right) in presence of 10 μM oligonucleotide T_20_ in standard conditions (top) and reducing conditions of 1 mM TCEP (bottom). Under non-reducing conditions, for equivalent concentrations of N-protein and T_20_, observed particles for N:G215C (**C**) are more fibrous than those for Nref (**A**). Such morphology can also be observed for Nref under slightly different conditions, and has been reported by others (Carlson et al., 2020). Addition of 1 mM TCEP as a reducing agent slightly accelerated droplet formation for Nref (**B**). Since Nref does not have any cysteines, this effect emphasizes the dependence of LLPS on protein solvation; for example, preferential exclusion of co-solutes from the vicinity of the protein will tend to stabilize the dense phase. For N:G215C the presence of 1 mM TCEP leads to reduction of disulfide bonds and altered quaternary structure. In the presence of T_20_ this promotes the formation of slightly larger droplets for reduced N:G215C (**D**) than for Nref (**B**).

## References

1. McDonald I, Murray SM, Reynolds CJ, Altmann DM, Boyton RJ (2021) Comparative systematic review and meta-analysis of reactogenicity, immunogenicity and efficacy of vaccines against SARS-CoV-2. npj Vaccines 6(1). doi:10.1038/s41541-021-00336-1.

2. Li D, Sempowski GD, Saunders KO, Acharya P, Haynes BF (2022) SARS-CoV-2 Neutralizing Antibodies for COVID-19 Prevention and Treatment. Annu Rev Med 73(1):1–16.

3. Cox RM, Wolf JD, Plemper RK (2021) Therapeutically administered ribonucleoside analogue MK-4482/EIDD-2801 blocks SARS-CoV-2 transmission in ferrets. Nat Microbiol 6(1):11–18.

4. Pruijssers AJ, et al. (2020) Remdesivir Inhibits SARS-CoV-2 in Human Lung Cells and Chimeric SARS-CoV Expressing the SARS-CoV-2 RNA Polymerase in Mice. Cell Rep 32(3). doi:10.1016/j.celrep.2020.107940.

5. Xiong M, et al. (2021) What coronavirus 3C-like protease tells us: From structure, substrate selectivity, to inhibitor design. Med Res Rev 41(4):1965–1998.

6. Zhao M, et al. (2021) GCG inhibits SARS-CoV-2 replication by disrupting the liquid phase condensation of its nucleocapsid protein. Nat Commun 12(1):2114.

7. Planas D, et al. (2021) Reduced sensitivity of SARS-CoV-2 variant Delta to antibody neutralization. Nature 596(7871):276–280.

8. Syed AM, et al. (2021) Rapid assessment of SARS-CoV-2 evolved variants using virus-like particles. Science (80-) 6184:2021.08.05.455082.

9. Syed AM, et al. (2022) Omicron mutations enhance infectivity and reduce antibody neutralization of SARS-CoV-2 virus-like particles. medRxiv Prepr Serv Heal Sci. doi:10.1101/2021.12.20.21268048.

10. Mourier T, et al. (2022) SARS-CoV-2 genomes from Saudi Arabia implicate nucleocapsid mutations in host response and increased viral load. Nat Commun 13(1). doi:10.1038/s41467-022-28287-8.

11. Elbe S, Buckland-Merrett G (2017) Data, disease and diplomacy: GISAID’s innovative contribution to global health. Glob Challenges 1(1):33–46.

12. Munis AM, Andersson M, Mobbs A, Hyde SC, Gill DR (2021) Genomic diversity of SARS-CoV-2 in Oxford during United Kingdom’s first national lockdown. Sci Rep 11(1):1–10.

13. Hadfield J, et al. (2018) NextStrain: Real-time tracking of pathogen evolution. Bioinformatics 34(23):4121–4123.

14. Ye Q, West AMV, Silletti S, Corbett KD (2020) Architecture and self-assembly of the SARS-CoV-2 nucleocapsid protein. Protein Sci 29(9):1890–1901.

15. Rochman ND, et al. (2021) Ongoing global and regional adaptive evolution of SARS-CoV-2. Proc Natl Acad Sci U S A 118(29):1–10.

16. Popa A, et al. (2020) Genomic epidemiology of superspreading events in Austria reveals mutational dynamics and transmission properties of SARS-CoV-2. Sci Transl Med 12(573):1–14.

17. Holland JJ, Domingo E (1997) RNA virus mutations and fitness for survival. Annu Rev Microbiol 51:151–178.

18. Graudenzi A, Maspero D, Angaroni F, Piazza R, Ramazzotti D (2021) Mutational signatures and heterogeneous host response revealed via large-scale characterization of SARS-CoV-2 genomic diversity. iScience 24(2):102116.

19. Domingo E, Perales C (2019) Viral quasispecies. PLoS Genet 15(10):1–20.

20. Domingo E, Garcia-Crespo C, Perales C (2021) Historical Perspective on the Discovery of the Quasispecies Concept. Annu Rev Virol 8:51–72.

21. Acevedo A, Brodsky L, Andino R (2014) Mutational and fitness landscapes of an RNA virus revealed through population sequencing. Nature 505(7485):686–690.

22. Wylie CS, Shakhnovich EI (2011) A biophysical protein folding model accounts for most mutational fitness effects in viruses. Proc Natl Acad Sci U S A 108(24):9916–9921.

23. Braun KM, et al. (2021) Acute SARS-CoV-2 infections harbor limited within-host diversity and transmit via tight transmission bottlenecks. PLoS Pathog 17(8):1–26.

24. Chertow D, et al. (2021) SARS-CoV-2 infection and persistence throughout the human body and brain National Institutes of Health. Res Sq. doi:10.21203/rs.3.rs-1139035/v1.

25. Tokuriki N, Oldfield CJ, Uversky VN, Berezovsky IN, Tawfik DS (2009) Do viral proteins possess unique biophysical features? Trends Biochem Sci 34(2):53–59.

26. Starr TN, et al. (2020) Deep Mutational Scanning of SARS-CoV-2 Receptor Binding Domain Reveals Constraints on Folding and ACE2 Binding. Cell 182(5):1295-1310.e20.

27. Garvin MR, et al. (2021) Rapid expansion of SARS-CoV-2 variants of concern is a result of adaptive epistasis. BioRxiv:1–54.

28. Klein S, et al. (2020) SARS-CoV-2 structure and replication characterized by in situ cryo-electron tomography. Nat Commun 11(1):5885.

29. Yao H, et al. (2020) Molecular Architecture of the SARS-CoV-2 Virus. Cell 183(3):730–738.e13.

30. Tao K, et al. (2021) The biological and clinical significance of emerging SARS-CoV-2 variants. Nat Rev Genet 0123456789. doi:10.1038/s41576-021-00408-x.

31. Li B, et al. (2021) Viral infection and transmission in a large, well-traced outbreak caused by the SARS-CoV-2 Delta variant. medRxiv:2021.07.07.21260122.

32. Teyssou E, et al. (2021) The Delta SARS-CoV-2 variant has a higher viral load than the Beta and the historical variants in nasopharyngeal samples from newly diagnosed COVID-19 patients. J Infect 83(4):e1–e3.

33. Eyre DW, et al. (2021) The impact of SARS-CoV-2 vaccination on Alpha and Delta variant transmission. medRxiv:2021.09.28.21264260.

34. Stern A, et al. (2021) The unique evolutionary dynamics of the SARS-CoV-2 Delta variant Israel Consortium of SARS-CoV-2 sequencing. medRxiv:2021.08.05.21261642.

35. Marchitelli V, et al. (2021) Evidence for the dependence of the SARS-Cov-2 Delta high diffusivity on the associated N : G215C nucleocapsid mutation. Res Sq:1–9.

36. Chang CK, Hou MH, Chang CF, Hsiao CD, Huang TH (2014) The SARS coronavirus nucleocapsid protein - Forms and functions. Antiviral Res 103(1):39–50.

37. Zhao H, et al. (2021) Energetic and structural features of SARS-CoV-2 N-protein co-assemblies with nucleic acids. iScience 24(6):102523.

38. Cubuk J, et al. (2021) The SARS-CoV-2 nucleocapsid protein is dynamic, disordered, and phase separates with RNA. Nat Commun 12(1):1936.

39. Lu S, et al. (2021) The SARS-CoV-2 nucleocapsid phosphoprotein forms mutually exclusive condensates with RNA and the membrane-associated M protein. Nat Commun 12(1):502.

40. Carlson CR, et al. (2020) Phosphoregulation of Phase Separation by the SARS-CoV-2 N Protein Suggests a Biophysical Basis for its Dual Functions. Mol Cell 80(6):1092–1103.e4.

41. Iserman C, et al. (2020) Genomic RNA Elements Drive Phase Separation of the SARS-CoV-2 Nucleocapsid. Mol Cell 80:1078–1091.

42. Masters PS (2019) Coronavirus genomic RNA packaging. Virology 537(August):198–207.

43. Dinesh DC, et al. (2020) Structural basis of RNA recognition by the SARS-CoV-2 nucleocapsid phosphoprotein. PLOS Pathog 16(12):e1009100.

44. Zinzula L, et al. (2021) High-resolution structure and biophysical characterization of the nucleocapsid phosphoprotein dimerization domain from the Covid-19 severe acute respiratory syndrome coronavirus 2. Biochem Biophys Res Commun 538:54–62.

45. Gussow AB, et al. (2020) Genomic determinants of pathogenicity in SARS-CoV-2 and other human coronaviruses. Proc Natl Acad Sci 117(26):202008176.

46. Poran A, et al. (2020) Sequence-based prediction of SARS-CoV-2 vaccine targets using a mass spectrometry-based bioinformatics predictor identifies immunogenic T cell epitopes. Genome Med 12(1):1–15.

47. Koetzner CA, Hurst-Hess KR, Kuo L, Masters PS (2022) Analysis of a crucial interaction between the coronavirus nucleocapsid protein and the major membrane-bound subunit of the viral replicase-transcriptase complex. Virology 567:1–14.

48. Wu CH, et al. (2009) Glycogen synthase kinase-3 regulates the phosphorylation of severe acute respiratory syndrome coronavirus mucleocapsid protein and viral replication. J Biol Chem 284(8):5229–5239.

49. Bouhaddou M, et al. (2020) The Global Phosphorylation Landscape of SARS-CoV-2 Infection. Cell 182(3):685–712.e19.

50. Tugaeva K V., et al. (2021) The Mechanism of SARS-CoV-2 Nucleocapsid Protein Recognition by the Human 14-3-3 Proteins. J Mol Biol 433(8):166875.

51. Quayum ST, Hasan S (2021) Analysing the impact of the two most common SARS-CoV-2 nucleocapsid protein variants on interactions with membrane protein in silico. J Genet Eng Biotechnol 19(1):138.

52. Azad GK (2021) The molecular assessment of SARS-CoV-2 Nucleocapsid Phosphoprotein variants among Indian isolates. Heliyon 7(September 2020):e06167.

53. Schuck P, Zhao H (2017) Sedimentation Velocity Analytical Ultracentrifugation: Interacting Systems (CRC Press, Boca Raton, FL).

54. Alberti S, Dormann D (2019) Liquid–Liquid Phase Separation in Disease. Annu Rev Genet 53(1):annurev-genet-112618-043527.

55. Kim TH, et al. (2021) Interaction hot spots for phase separation revealed by NMR studies of a CAPRIN1 condensed phase. Proc Natl Acad Sci U S A 118(23):1–11.

56. Lin Y-H, Wu H, Jia B, Zhang M, Chan HS (2021) Assembly of model postsynaptic densities involves interactions auxiliary to stoichiometric binding. Biophys J in press. doi:10.1016/j.bpj.2021.10.008.

57. Wu C, et al. (2021) Characterization of SARS-CoV-2 nucleocapsid protein reveals multiple functional consequences of the C-terminal domain. iScience 24(6):102681.

58. Peng Y, et al. (2020) Structures of the SARS-CoV-2 nucleocapsid and their perspectives for drug design. EMBO J 39(20):e105938.

59. Zhou R, Zeng R, von Brunn A, Lei J (2020) Structural characterization of the C-terminal domain of SARS-CoV-2 nucleocapsid protein. Mol Biomed 1(1):2.

60. Schiavina M, Pontoriero L, Uversky VN, Felli IC, Pierattelli R (2021) The highly flexible disordered regions of the SARS-CoV-2 nucleocapsid N protein within the 1–248 residue construct: sequence-specific resonance assignments through NMR. Biomol NMR Assign 15(1):219–227.

61. Rubayet Ul Alam ASM, et al. (2021) Dominant Clade-featured SARS-CoV-2 Co-occurring Mutations Reveals Plausible Epistasis: An in silico based Hypothetical Model. J Med Virol. doi:10.1002/jmv.27416.

62. Del Veliz S, Rivera L, Bustos DM, Uhart M (2021) Analysis of SARS-CoV-2 nucleocapsid phosphoprotein N variations in the binding site to human 14-3-3 proteins. Biochem Biophys Res Commun 569:154–160.

63. Savastano A, Ibáñez de Opakua A, Rankovic M, Zweckstetter M (2020) Nucleocapsid protein of SARS-CoV-2 phase separates into RNA-rich polymerase-containing condensates. Nat Commun 11(1):6041.

64. Haibo Wu, Na Xing, Kaiwen Meng, Beibei Fu, Weiwei Xue, Pan Dong, Yang Xiao, Gexin Liu, Haitao Luo, Wenzhuang Zhu, Xiaoyuan Lin, Geng Meng ZZ (2021) Nucleocapsid mutation R203K/G204R increases the infectivity, fitness and virulence of SARS-CoV-2. BioRxiv (55). doi:10.1101/2021.05.24.445386.

65. Bessa LM, et al. (2022) The intrinsically disordered SARS-CoV-2 nucleoprotein in dynamic complex with its viral partner nsp3a. Sci Adv 8(3). doi:10.1126/sciadv.abm4034.

66. Wootton SK, Yoo D (2003) Homo-Oligomerization of the Porcine Reproductive and Respiratory Syndrome Virus Nucleocapsid Protein and the Role of Disulfide Linkages. J Virol 77(8):4546–4557.

67. Prokudina EN, Semenova NP, Chumakov VM, Rudneva IA (2004) Transient disulfide bonds formation in conformational maturation of influenza virus nucleocapsid protein (NP). Virus Res 99(2):169–175.

68. Wu F, et al. (2020) A new coronavirus associated with human respiratory disease in China. Nature 579(7798):265–269.

69. Papadopoulos JS, Agarwala R (2007) COBALT: constraint-based alignment tool for multiple protein sequences. Bioinformatics 23(9):1073–9.

70. Robert X, Gouet P (2014) Deciphering key features in protein structures with the new ENDscript server. Nucleic Acids Res 42(W1):W320–W324.

71. Rao JKM (2009) New scoring matrix for amino acid residue exchanges based on residue characteristic physical parameters. Int J Pept Protein Res 29(2):276–281.

72. Hardenberg M, Horvath A, Ambrus V, Fuxreiter M, Vendruscolo M (2020) Widespread occurrence of the droplet state of proteins in the human proteome. Proc Natl Acad Sci:202007670.

73. Jumper J, et al. (2021) Highly accurate protein structure prediction with AlphaFold. Nature 596(7873):583–589.

74. Yang J, et al. (2015) The I-TASSER Suite: protein structure and function prediction. Nat Methods 12(1):7–8.

75. Kelley LA, Mezulis S, Yates CM, Wass MN, Sternberg MJE (2015) The Phyre2 web portal for protein modeling, prediction and analysis. Nat Protoc 10(6):845–858.

76. Brooks BR, et al. (2009) CHARMM: The biomolecular simulation program. J Comput Chem 30(10):1545–1614.

77. Huang J, et al. (2017) CHARMM36m: an improved force field for folded and intrinsically disordered proteins. Nat Methods 14(1):71–73.

78. Schuck P, Zhao H, Brautigam CA, Ghirlando R (2015) Basic Principles of Analytical Ultracentrifugation (CRC Press, Boca Raton, FL).

79. Schuck P (2016) Sedimentation Velocity Analytical Ultracentrifugation: Discrete Species and Size-Distributions of Macromolecules and Particles (CRC Press, Boca Raton, FL).

80. Wu D, Piszczek G (2021) Standard protocol for mass photometry experiments. Eur Biophys J 50(3-4):403–409.

